# Vascular damage and excessive proliferation compromise liver function after extended hepatectomy in mice

**DOI:** 10.1101/2024.01.04.573041

**Authors:** Maxime De Rudder, Rita Manco, Laurent Coubeau, Alix Fontaine, Claude Bertrand, Isabelle A. Leclercq, Alexandra Dili

## Abstract

Surgical resection remains the gold standard for liver tumor treatment, yet the emergence of post-operative liver failure, known as the small for size syndrome (SFSS), poses a substantial challenge. The activation of hypoxia sensors in a SFSS liver remnant initiated early angiogenesis, improving vascular architecture, safeguarding against liver failure and reducing mortality. The study aimed to elucidate vascular remodeling mechanisms in SFSS, its impact on hepatocyte function and subsequent liver failure. Mice underwent extended partial hepatectomy to induce SFSS, a subset were exposed to hypoxia immediately after surgery. Hypoxia bolstered post-hepatectomy survival rates. Early proliferation of liver sinusoidal cells coupled with augmented recruitment of endothelial progenitor cells (EPC) via the VEGF/SDF-1α pathway resulted in heightened vascular density, improved lobular perfusion, and limited hemorrhagic events in the regenerating liver under hypoxia. The administration of G-CSF mimicked the effects of hypoxia on vascular remodeling and EPC recruitment, though it failed to rescue survival. Compared to normoxia, hypoxia restrained hepatocyte proliferation yet improved the function of the regenerating remnant, favoring functional preservation in the liver remnant. Injection of AAV8- TBG-HNF4α virus for hepatocyte-specific overexpression of HNF4α, the master regulator of hepatocyte function, enforced functionality in proliferating hepatocytes. The combination, only, of HNF4α overexpression and G-CSF treatment rescued survival post-SFSS-setting hepatectomy. In summary, SFSS arises due to imbalance and desynchronized interplay between functional regeneration and vascular restructuring. To enhance survival following SFSS-hepatectomy, a two-pronged strategy is essential, addressing the preservation of function in the proliferating parenchymal cells alongside the simultaneous mitigation of vascular harm.

**One Sentence Summary:** Combined treatment with G-CSF and HNF4α overexpression rescues vascular damage and function to improve survival after extended hepatectomy in mice.

## INTRODUCTION

Surgical resection is the preferred treatment for primary or secondary liver tumor, offering the patient the best chance for long-term survival (*1, 2*). However, despite preconditioning strategies and improvement in surgical techniques (*1, 3–7*), there is still a risk of post-operative liver failure, also known as the small for size syndrome (SFSS) (*8*). The incidence of SFSS ranges between 1.2 and 35%, with a mortality rate of up to 70% (*9*). The SFSS, also observed after transplantation of small liver grafts (*10*), significantly reduces the number of patients eligible for a potentially curative treatment. SFSS diagnosis is typically based on clinical and biological features on postoperative day 5 or day 14 (*10, 11*), when irreversible parenchymal damage has already occurred, leaving little opportunity for treatment.

The main factor that contributes to liver insufficiency is the increased portal perfusion exceeding four times the baseline which triggers a proportionate hepatic arterial buffer response (*12,13*). The increased shear stress leads to focal disruption of the sinusoidal endothelial lining, endothelial denudation, connective tissue hemorrhage and edema in early liver histopathological findings of SFSS livers (*14*). Other observed features include sinusoidal congestion, irregular large gaps of sinusoidal lining cells, collapse of the space of Disse and hepatocyte ballooning. Simultaneously, the hepatic arterial buffer response (*12*) causes constriction of the hepatic artery and leads to de- arterialization, and ischemic injury (*15–17*). As such, histopathological observations show arterial vasospasm and thrombosis, perihilar bile duct necrosis, cholangitis abscesses and parenchymal infracts at later stages of SFSS after liver transplantation (10–20 days) (*14, 18*). Portal hyperperfusion, de-arterialization and the resulting hypoxia in the remnant liver are thus proposed as primary factors for life-threatening postoperative liver failure, which prompts some authors to rename this entity as the *small-for-flow-syndrome* (*19*).

While disturbances in the inflow of the liver remnant are commonly regarded as the primary factors causing SFSS, studies in both animals and humans have demonstrated that increased portal perfusion is necessary for liver regeneration. The magnitude of the regenerative stimulus is proportional to the rise in portal blood flow (*20–23*). Though, a high regenerative response, seen as an extreme hepatocyte proliferation rate, is associated with the formation of avascular hepatocyte clusters, disruption of the lobular architecture and a higher risk of liver insufficiency after extended hepatectomy or transplantation of a small graft (*22*).

Thus, portal overflow, disturbance in hepatic microcirculation and lobular disorganization along with hepatocyte hyperproliferation concur to hepatocyte dysfunction and SFSS (*24*). We already demonstrated that the activation of hypoxia sensors in a SFSS liver remnant initiates an early angiogenic switch. This switch improved vascular architecture in the regenerating organ and prevented mortality (*25*). To the best of our knowledge, it is currently unclear whether dysfunctional liver regeneration after extended hepatectomy is linked to a vascular problem leading to hepatocyte dysfunction, or whether hepatocytes are incapable to face the metabolic load of the small liver remnant, regardless of the status of the sinusoidal network. To follow-up on this, the primary aim of our study is to investigate the mechanisms behind vascular remodeling in a small liver remnant. Additionally, we aim to determine how this remodeling impacts hepatocyte function and whether it is sufficient to prevent liver failure and SFSS.

## RESULTS

### Hypoxia improves lobular perfusion and the sinusoidal vascular network in a SFS remnant

Eighty percent partial hepatectomy (PHx80%) is a model of SFSS in mice as it causes 7-day mortality in 60% of animals. Most of lethal events occurred between the first and third day after surgery, suggesting that liver dysfunction occurs in the early phase of liver regeneration. Based on this observation, we focused our research on that early timing for experimental manipulation. When mice undergoing extended hepatectomy were exposed to hypoxia immediately after surgery and for 3 consecutive days (PHx80%-HC), the survival significantly improved, resulting in a reduction of the seven-day mortality from 61.8% to 31.8% (p=0.003) (Fig 1A).

**Fig. 1.**
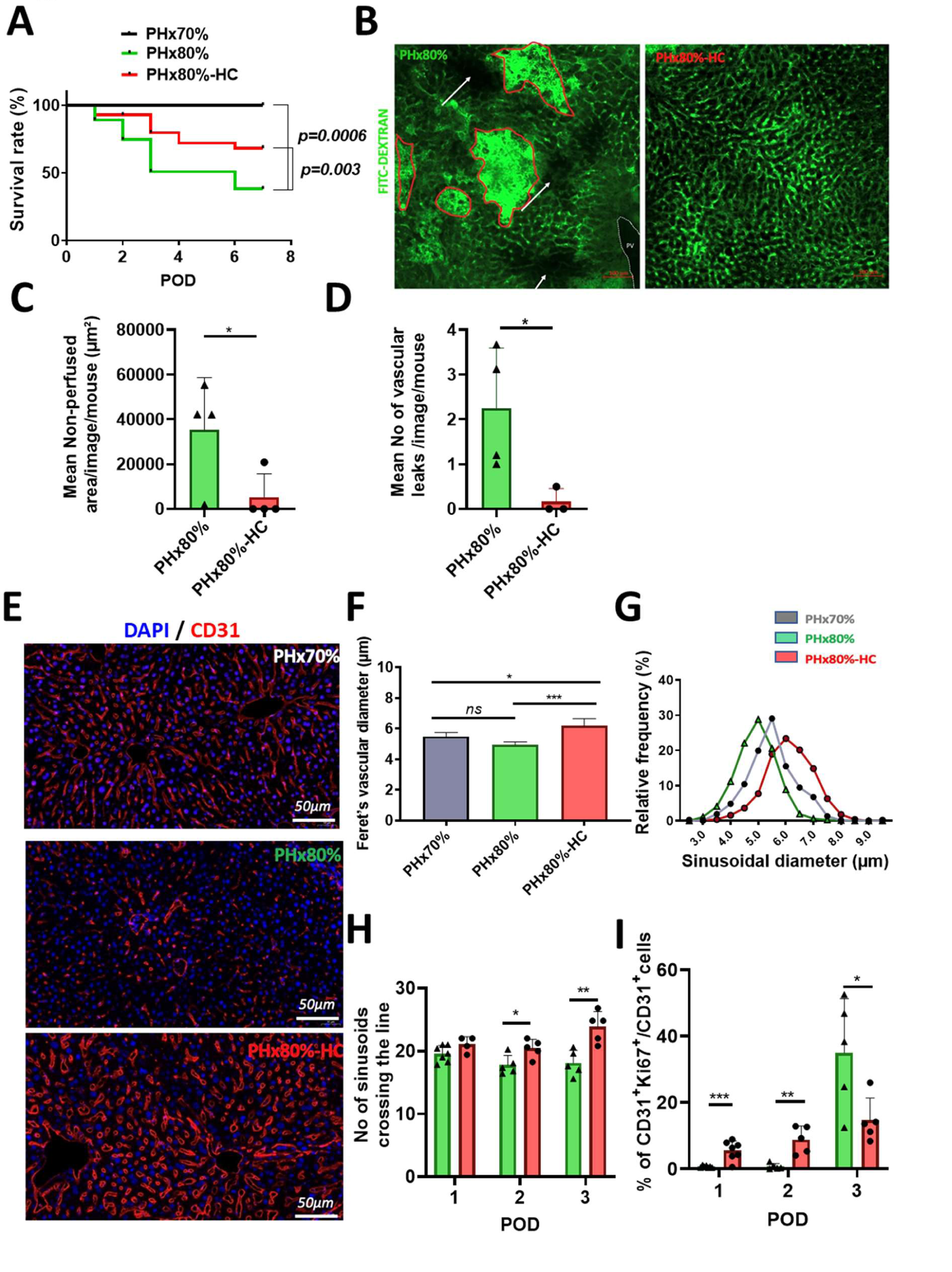
Hypoxia increases survival after a SFSS-setting hepatectomy, improves lobular perfusion, prevents vascular leaks and contributes to the vascular remodeling in the rapidly regenerating liver. (A) Kaplan Meier survival curves after PHx70% (Dark grey line; n=19), PHx80% (or SFSS-setting hepatectomy) (green line; n=73) and PHx80%-HC (red line; n=56) in C57Bl6/J mice. Log-rank (Mantel-Cox) test was used as statistical test. **(B-D)** Liver perfused with fixable FITC-Dextran in PHx80% and PHx80%-HC on POD3. **(B)** Representative picture of liver sections; w*hite arrows* indicate dark, non-perfused areas, while *red lines* show vascular leaks with intraparenchymal collection of the intravascular fluorescent green dye. PV: portal vein. (Bar size = 100µm). **(C)** Quantification of the non-perfused area in the regenerating liver in PHx80% (n=4) and PHx80%-HC (n=4) on POD3. *(*p=0.05)* and of **(D)** the number of vascular leaks in the regenerating liver in PHx80% (n=4) and PHx80%-HC (n=3) on POD3. *(*p=0.04)*. **(E)** Representative pictures of sinusoids stained with CD31 in the regenerating liver in PHx80% and PHx80%-HC on POD1. (Bar size = 50µm). **(F)** Quantification of the mean sinusoidal diameter in PHx70% (n=4), PHx80% (n=6), PHx80%-HC (n=4) on POD1 and **(G)** distribution of the sinusoidal diameter in PHx70% (n=4), PHx80% (n=6), PHx80%-HC (n=4) on POD1 (\*\*\**p=0.0005 in* PHx80% vs PHx80%-HC). **(H)** Sinusoidal density in PHx80% at POD1 (n=7), POD2 (n=5) and POD3 (n=5), and PHx80%-HC at POD1 (n=4), POD2 (n=5) and POD3 (n=5). unpaired Multiple t-test (POD2: *(*p=0.02)*, POD3 *(**p=0.003)*). **(I)** Liver sinusoidal endothelial cell proliferation in liver remnants after SFSS-setting hepatectomy in PHx80% at POD1 (n=9), POD2 (n=5) and POD3 (n=5), and PHx80%-HC at POD1 (n=7), POD2 (n=5) and POD3 (n=5). Multiple unpaired t-test (POD1: *(***p=0.0007),* POD2: *(**p=0.002)*, POD3 *(*p=0.03)*).

To assess liver microperfusion, we injected fixable fluorescein-Dextran, a high molecular weight intravascular fluorescent dye, three days after extended partial hepatectomy in mice kept or not in hypoxia chamber during the postoperative period. After PHx80%, liver remnants presented inadequate perfusion, which was indicated by the presence of multiple non-perfused regions in alternance with area of vascular leakages and extravasation of the fluorescent dye (Fig 1B). In contrast, immersion of the animal into a hypoxic environment significantly improved the lobular perfusion (p= 0.05) (Fig 1C) and limited hemorrhagic events (p=0.04) (Fig 1D) in the regenerating liver remnant despite undergoing the same extent of liver resection.

We examined liver sinusoids with CD31 IF staining of liver sections. In normoxic post-PHx80% remnants, hepatic sinusoids were small or even collapsed, whereas hypoxia shaped a network of large and patent sinusoidal vessels alike the one observed after a well-tolerated PHx70% (Fig 1E). Morphometric quantifications confirmed the difference. Compared to PHx80%, the mean diameter of sinusoid vessels in the liver remnant was significantly larger already at POD1 (p=0.0005) (Fig 1F), with the distribution of vessels’ caliber shifting towards larger sinusoids (Fig 1G) in mice housed in the hypoxia chamber. The vascular density was significantly higher at POD2 (p=0.02) and POD3 as well (p=0.003) (Fig 1H). Quantification of the proliferative endothelial cells showed that hypoxia triggered the proliferation of liver sinusoidal endothelial cells (LSEC) already at POD1 (p=0.0007) and POD2 (p=0.002) (Fig 1I), while LSEC proliferation in normoxic environment was not detected before POD3 (p=0.03). Taken together, our data support that hypoxia improves the lobular perfusion in a context of SFSS-setting hepatectomy through early endothelial proliferation and vascular remodeling.

### Hypoxia after SFSS hepatectomy favors liver function and lobular organization over liver mass restitution

Additionally, we observed that exposure to a hypoxic environment improved the function of the small remnant. At postoperative day 3, serum albumin and factor V were significantly higher in the PHx80%-HC group compared to the normoxic group (p=0.004 and p<0.0001 respectively) (Fig 2A+B). This indicates improved liver function in the hypoxic group. Consistently, mRNA levels of genes involved in the overall hepatocyte function were significantly increased by hypoxia, notably *Cyp2e1*, *Cyp1a2*, *Glul*, *Apob*, *F5* and *F12* (Fig 2C). Intracellular glycogen storage serves as an indicator of hepatocyte glycogen synthetic capacity and hepatocyte function. After extended hepatectomy, and compared to 70% PHx, significant depletion of glycogen was observed in many hepatocytes (PHx70% vs PHx80%, p=0.002). However, in mice exposed to hypoxia, the depletion was much less pronounced (p=0.2) and glycogenic hepatocytes were significantly more numerous compared to mice with extended hepatectomy in the normoxic group (p=0.03) indicating improved early functional recovery (Fig 2D+E).

**Fig. 2.**
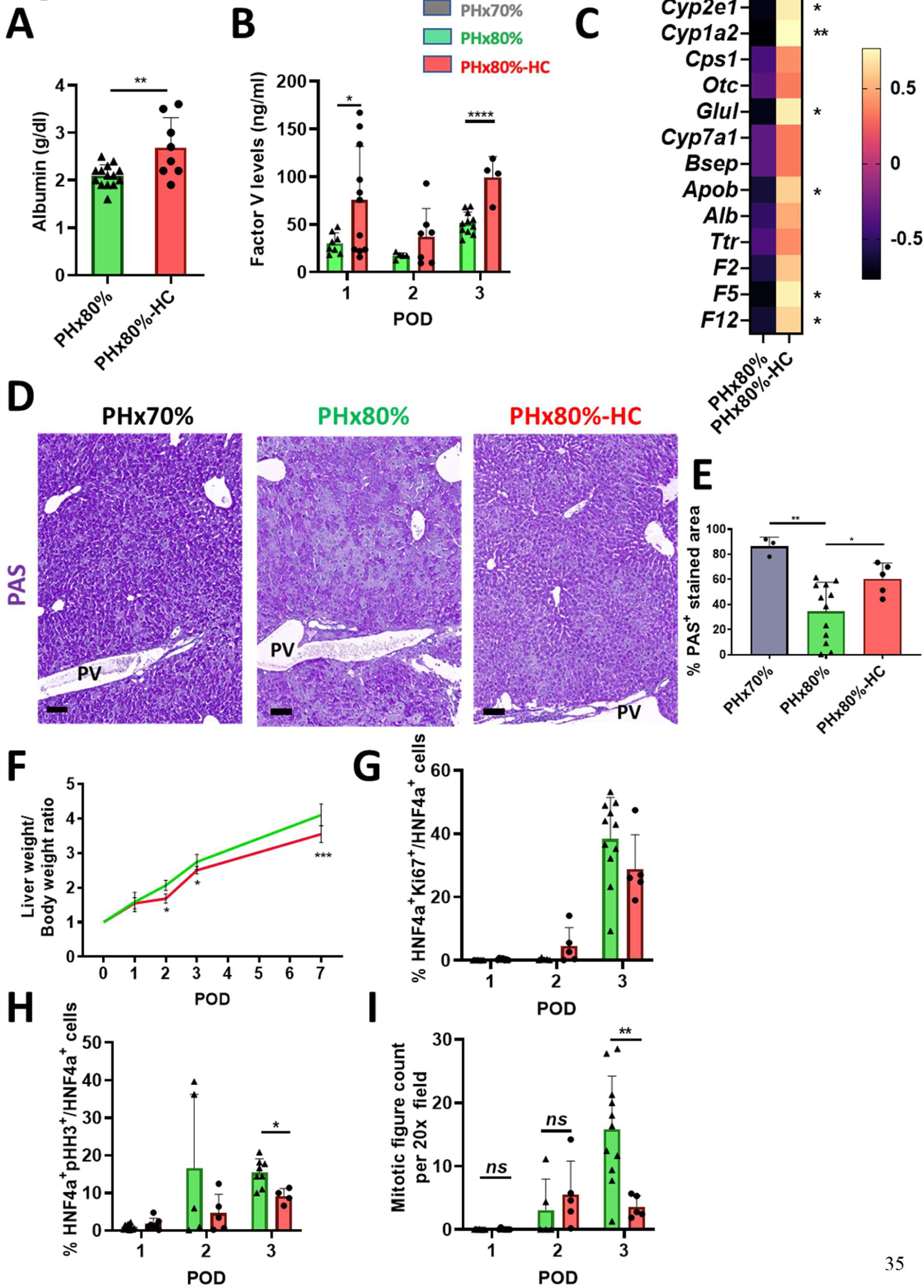
Hypoxia favors early functional liver recovery over liver mass recovery in the setting of SFSS hepatectomy. **(A)** Albumin plasma levels (g/dl) in SFSS-setting hepatectomy in PHx80% (n=14) and PHx80%- HC (n=8) on POD3. (\*\**p=0.004*). **(B)** Factor V plasma protein levels in PHx80% on POD1 (n=8), POD2 (n=5), POD3 (n=11) and in PHx80%-HC (n=11), POD2 (n=7), POD3 (n=4). Multiple unpaired t-test, POD1 *(*p=0.03)* and POD3 *(***p<0.0001).* **(C)** Heatmap of genes expressions reflecting hepatocyte function in PHx80% (n=5), PHx80%-HC (n=5) on POD3. Z-score was calculated from DeltaCT and plotted on the heatmap. **(D)** Representative pictures of PAS staining in the regenerative livers in PHx70%, PHx80% and PHx80%-HC on POD3 (Bar size = 100µm) and **(E)** quantification of the PAS-stained area/total slide area in PHx70% (n=3), PHx80% (n=12) and PHx80%-HC (n=5) on POD3. (PHx80% vs PHx80%-HC *(*p=0.03)*). **(F)** Liver remnant mass recovery expressed as liver remnant weight in relation to body weight in PHx80% and PHx80%- HC [PHx80% (n=18 on POD1, n=5 on POD2, n=17 on POD3, n=3 on POD7), PHx80%-HC (n=10 on POD1, n=5 on POD2, n=9 on POD3, n=11 on POD7)]. Two-way ANOVA (POD2 \**p=0.02*; POD3 \**p=0.02*; POD7 \*\*\**p=0.0005*). **(G)** Quantification of Ki67 positive hepatocytes during the early phase of post-hepatectomy liver regeneration in PHx80% on POD1 (n=8), POD2 (n=5) and POD3 (n=11) and PHx80%-HC on POD1 (n=7), POD2 (n=5) and POD3(n=5). **(H)** Quantification of pHH3 positive hepatocytes in PHx80% on POD1 (n=11), POD2 (n=5) and POD3 (n=8), and PHx80%-HC on POD1 (n=7), POD2 (n=5) and POD3(n=4) during the early phase of post- hepatectomy liver regeneration. Multiple t-test unpaired analysis (POD3*, (*p=0.01).* **(I)** Quantification of mitotic features in 20x/field in PHx80% on POD1 (n=8), POD2 (n=5) and POD3 (n=11), and PHx80%-HC on POD1 (n=7), POD2 (n=5) and POD3 (n=5) during the early phase of post-hepatectomy liver regeneration. Multiple t-test unpaired analysis *(**p=0.007*).

Surprisingly, the immediate functional recovery contrasted with poor restitution of the liver remnant mass. From postoperative day 2, liver mass recovery was significantly lower in the SFSS- setting hepatectomy under hypoxic conditions compared to survivors of normoxic extended hepatectomy (Fig 2F). To understand this paradox, we extensively assessed parenchymal cell proliferation. Hepatocyte proliferation at postoperative day 1 and 2 showed no difference between groups whether based on Ki67, pHH3 expression or on the number of mitoses (Fig 2G+H+I). However, at day 3, there were significantly fewer mitotic hepatocyte features in the remnants exposed to hypoxia compared to normoxia (p=0.007) (Fig 2I). This was also confirmed by a significantly lower proportion of pHH3 positive hepatocyte nuclei (p=0.01) (Fig 2H). Because the peak of deaths and hepatocyte proliferation happens at POD3 in our model, we decided to mainly focus on that timepoint for further experiments.

Since liver function is influenced by the lobular architecture (*26*), we investigated whether the early functional recovery of the liver remnant under hypoxic conditions was associated with improved interconnection of hepatocytes to the remodeled sinusoidal bed, and thus an improved lobular architecture during organogenesis. We used E-cadherin to stain hepatocyte contours and CD31 to delineate LSEC (Fig 3A). We observed clusters of hepatocytes disconnected from the vascular bed in liver remnants after extended hepatectomy, while in mice exposed to hypoxia the hepatocyte avascular islets were significantly fewer during the early phase of regeneration (p=0.0004) (Fig 3A+B).

**Fig. 3.**
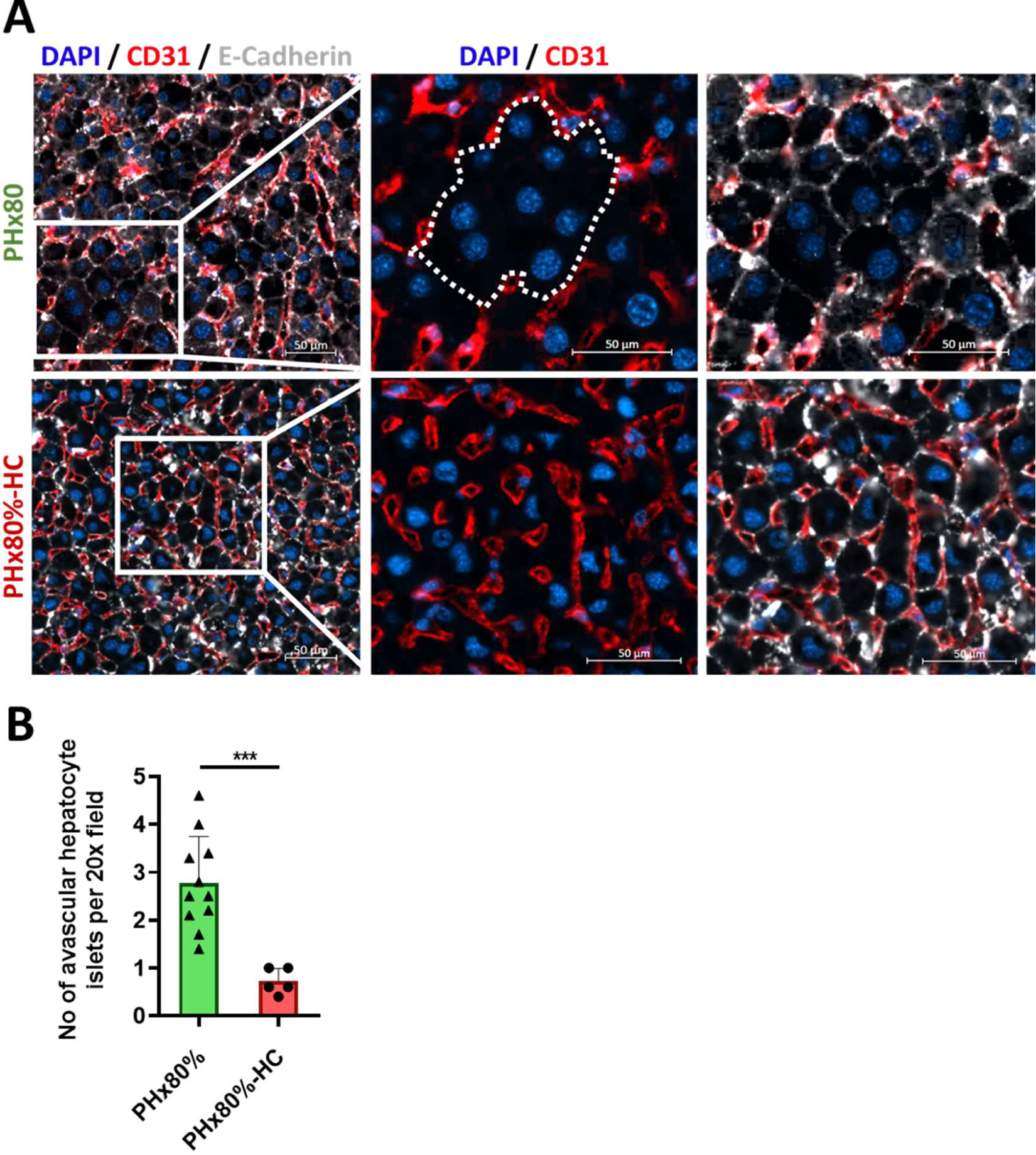
In the setting of SFSS hepatectomy, hypoxia prevents the formation of avascular hepatocyte islets. **(A)** Representative images of E-cadherin *(in white)*, CD31 *(in red)* and nucleus (Dapi *in blue*)IF staining in PHx80% and PHx80%-HC on POD3 (Bar size = 50µm) and **(B)** quantification of the number of hepatic clusters per image in PHx80% (n=12) and PHx80%-HC (n=5) on POD3. Unpaired t-test analysis (\*\*\**p=0.0004*).

Based on these findings, we propose that hypoxia induces early vascular regeneration coupled with reduced hepatocyte proliferation, facilitates the organization of hepatocytes in plates along sinusoids and enables functional compensation of the lost liver mass, hence, improving the animal’s survival.

### Hypoxia-driven recruitment of endothelial precursors contributes to angiogenesis in the SFSS liver remnant

Previous studies have demonstrated the recruitment of endothelial progenitor cells (EPC) to the liver in wound healing (*27*), endothelial cell X-ray damage (*28*), and partial hepatectomy (*29*), though the specific cellular source of regenerating endothelium varies according to the fitness of the residual vascular network (*30*). In our study, we showed that hypoxia rescued vascular damage after extended hepatectomy. To evaluate whether endothelial progenitors contribute to the vascular remodeling in the hypoxic SFSS remnant, we used Cdh5-PAC-Cre^ERT2^XROSA^mT/mG^ mice to permanently label with GFP endothelial cells and their progeny prior to surgery. Our labeling system achieved high efficiency, with over 90% of VE-Cadherin positive cells showing GFP label as confirmed by flow cytometry (Suppl Fig 1B+C) and immunofluorescence (Suppl Fig 2D) in control livers.

Three days after a SFSS-setting hepatectomy (PHx80%) the proportion of unlabeled endothelial cells increased to 20% of LSEC (Fig 4A+B). In the liver of mice exposed to hypoxia after PHx80%, we identified up to 35% of the endothelial cells that were not GFP labelled (Fig 4A+B). These findings support that, in hypoxic conditions, novel endothelial cell precursors contribute to vascular regeneration. The VEGF/SDF-1α pathway was highly activated in the hypoxic group, (Fig 4C+D), consistent with angiogenesis and the recruitment of endothelial progenitors (*31*). Similarly, levels of GM-CSF, another relevant factor for mobilization of bone marrow progenitors, were also increased in the plasma (Fig 4E).

**Fig. 4.**
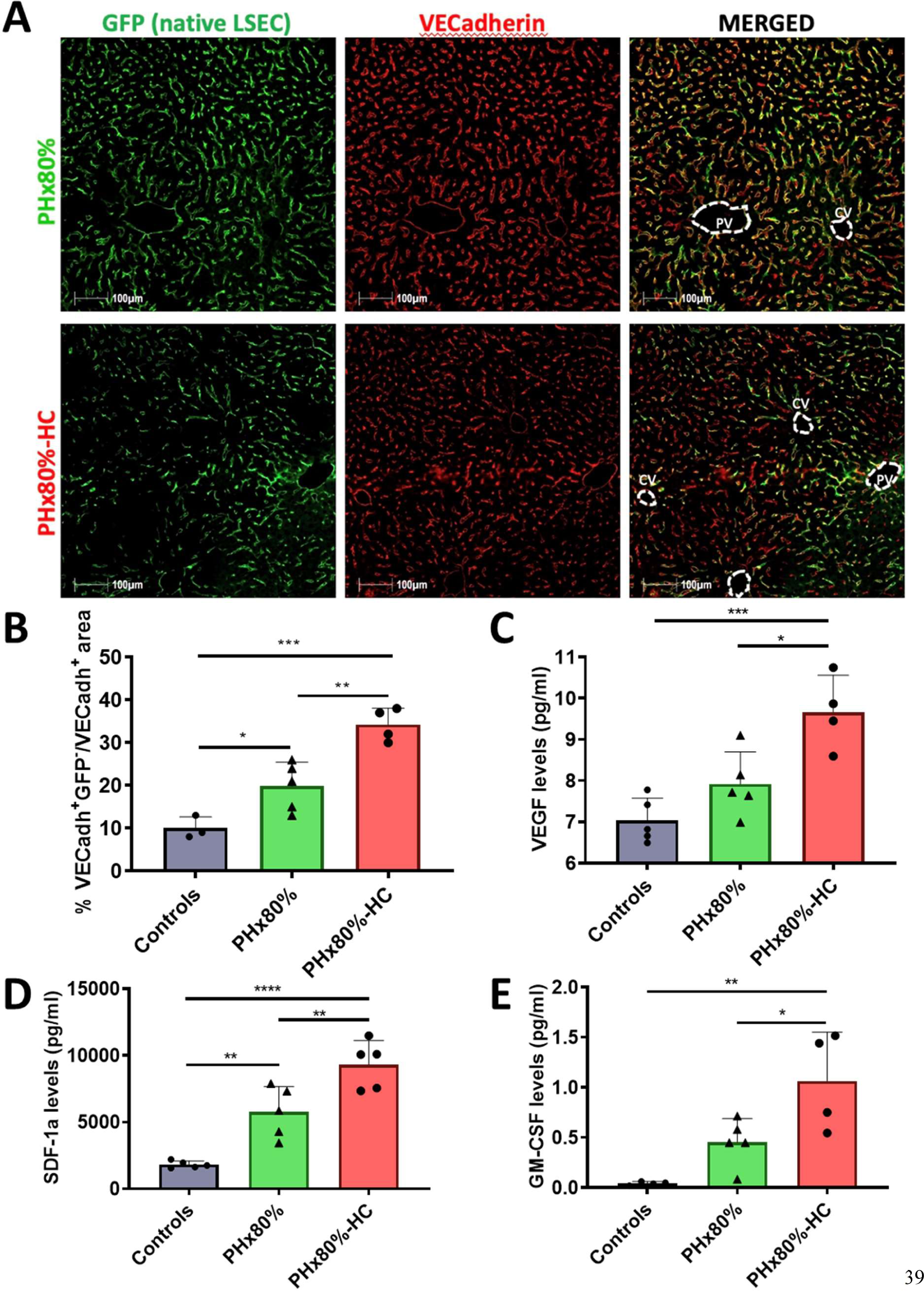
Hypoxia-driven recruitment of endothelial progenitors contributes to angiogenesis in the SFSS liver remnant. PHx80% and PHx80%-HC in Cdh5-PAC-Cre^ERT2^XROSA^mT/mG^ **(A)** Representative pictures of IF in the regenerating livers illustrating GFP^+^ cells (*in green,* liver native endothelial cells, left panels), VECadh^+^ cells (*in red*, all endothelial cells, middle panels) and merged images (right panels) in PHx80% and PHx80%-HC. CV: central vein, PV: portal vein. (Bar size = 100µm). **(B)** Quantification of the percentage of VECadh^+^GFP^-^ endothelial cells (non-liver native endothelial cells) in the regenerating liver in controls (Tamoxifen injected and untouched) (n=3), PHx80% (n=5) and PHx80%-HC (n=4) on POD3. (PHx80% vs PHx80%-HC, *\*\*p=0.003*). Plasma concentrations of **(C)** VEGF (PHx80% vs PHx80%-HC, *\*p=0.01*), **(D)** SDF-1α (PHx80% vs PHx80%-HC, *\*\*p=0.009)* and **(E)** GM-CSF (PH80% vs PH80%-HC, \**p=0.04*). in controls (n=5), PHx80% (n=5) and PHx80%-HC (n=4 or 5) on POD1. Data are analyzed by One-way ANOVA.

### Mobilization of bone marrow progenitors mimics the beneficial effects of hypoxia-induced angiogenesis after extended hepatectomy

According to the literature, endothelial progenitors derived from the bone marrow have been shown to home in the regenerating liver after 70% partial hepatectomy (*32*). Therefore, we investigated the impact of mobilization of bone marrow-derived endothelial progenitor cells (BM-EPC) on early angiogenesis in the SFS remnant using G-CSF in Cdh5-CreER^T2^ X Rosa^mT/mG^ mice. G-CSF increased the number of circulating white blood cells without disturbing the other blood cell types (Suppl Fig 2A+B). In PHx80% mice, G-CSF increased the number of GFP^-^ endothelial cells recruited to the liver remnant compared to vehicle controls (p=0.002) (Fig 5A). This goes in parallel with increased vascular density and sinusoidal diameter in the regenerating liver remnants (p<0.0001 and p=0.01, respectively) (Fig 5B+C). Thus, mobilization of bone-marrow progenitors enhanced endothelial cell engraftment and preserved the vascular network in the regenerating liver as seen with hypoxia.

**Fig. 5.**
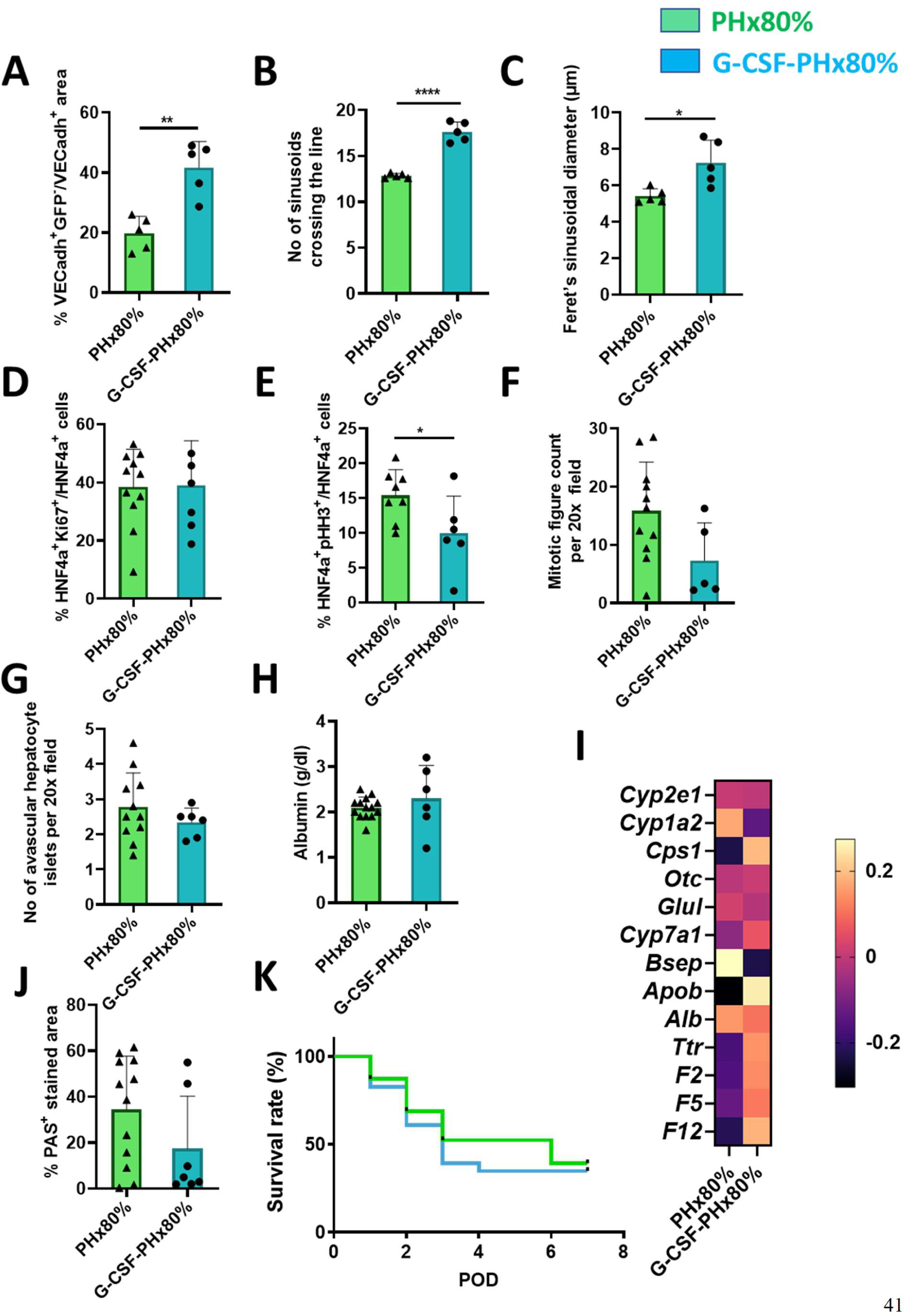
Recruitment of putative bone marrow progenitors mimics hypoxia-induced angiogenesis and vascular remodeling in Cdh5-Cre x mTmG but does not improve lobular architecture and function in C57Bl/6J. Cdh5-PAC-Cre^ERT2^XROSA^mT/mG^ mice underwent PHx80% with or without treatment with G-CSF. PHx80% (n=5) and G-CSF-PHx80% (n=5). Analysis at POD3 **(A)** Percentage of VECadh^+^GFP^-^ endothelial cells (non-liver native endothelial cells) (\*\**p=0.002).* **(B)** Sinusoidal density on CD31 IF stained sections *(****p<0,0001)* **(C)** Mean sinusoidal diameter (\**p=0.01*) C57Bl/6J mice underwent PHx80% with or without treatment with G-CSF. **(D)** Percentage of Ki67 positive HNF4α+ hepatocytes on POD3 in PHx80% (n=11), G-CSF-PHx80% (n=7) in C57Bl/6J mice. **(E)** Quantification of pHH3 positive hepatocytes in PHx80% (n=8) and G-CSF-PHx80% (n=6) on POD3. *(*p=0.04)* and **(F)** of mitotic features in 20x/field on POD3 in PHx80% (n=11) and G-CSF-PHx80% (n=5). **(G)** Quantification of the number of hepatic clusters on POD3 in PHx80% (n=12) and G-CSF-PHx80% (n=6). **(H)** Albumin plasma levels (g/dl) in SFSS-setting hepatectomy in PHx80% (n=14) and in G-CSF-PHx80% (n=6) on POD3. **(I)** Heatmap of genes expressions reflecting hepatocyte function on POD3. Z-score was calculated from DeltaCT and plotted on the heatmap for PHx80% (n=5), G-CSF-PHx80% (n=6) **(J)** Quantification of the PAS- stained area/total slide area in PHx80% (n=12) and G-CSF-PHx80% (n=7) on POD3. **(K)** Kaplan Meier survival curves after PHx80% (n=73) and G-CSF-PHx80% (n=23) in C57Bl6/J mice.

In contrast to reduced hepatocyte proliferation seen in the hypoxic group, G-CSF treated mice showed similar number of hepatocytes in the cell cycle (Fig 5D) or in mitosis as in vehicle controls (Fig 5E), even though pHH3^+^ hepatocytes were fewer (Fig 5F). The number of hepatocyte islets disconnected from the sinusoidal network was also similar (Fig 5G). Serum albumin levels (Fig 5H), expression of functional genes (Fig 5I), PAS staining (Fig 6J) as well as low survival rate (Fig 5K) all demonstrate that salvage of the vascular bed is not sufficient to support function during the regeneration after extended hepatectomy.

**Fig. 6.**
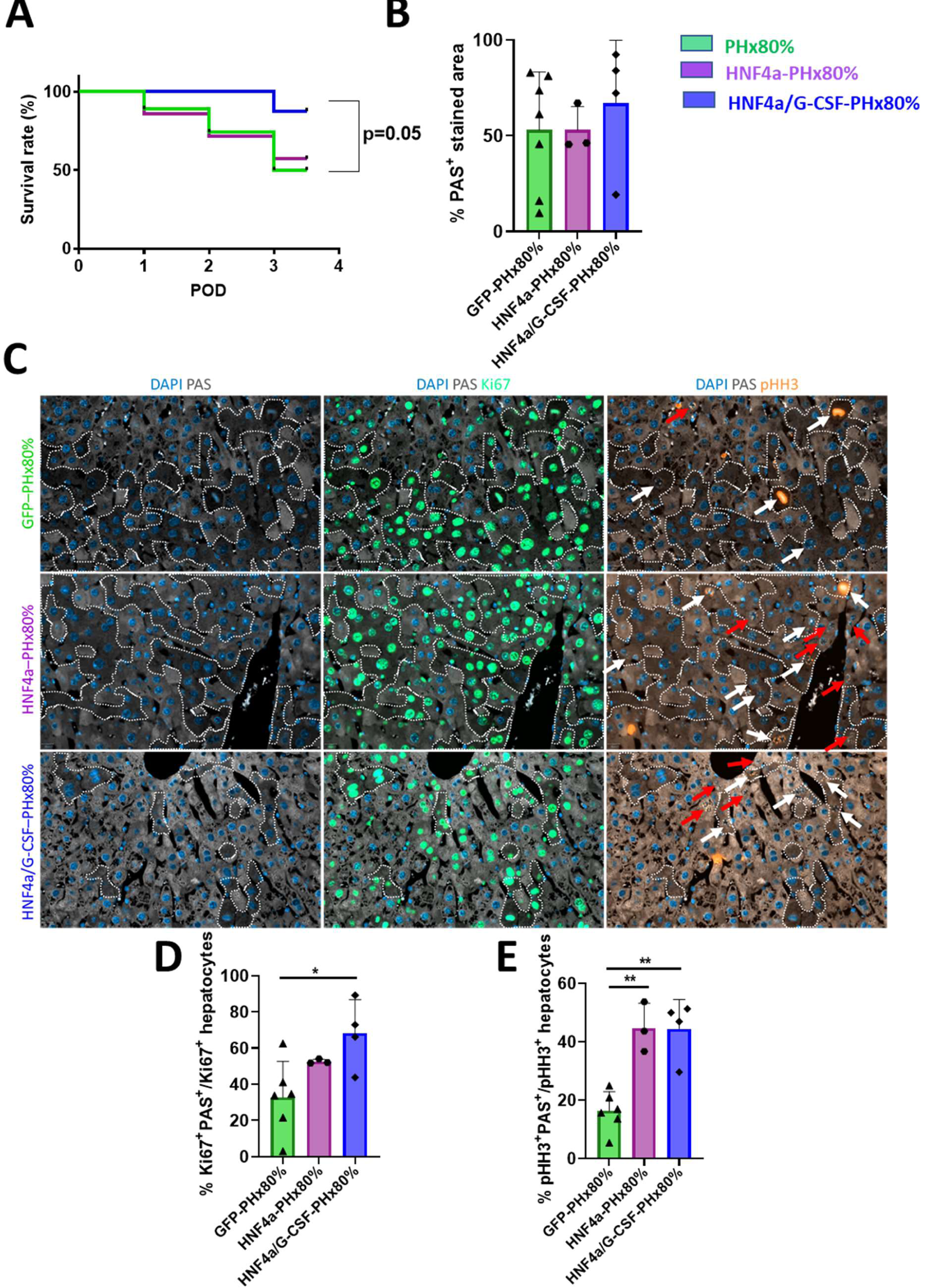
G-CSF treatment combined with HNF4α overexpression increases survival and induces a hybrid hepatocyte phenotype in mice after extended hepatectomy. **(A)** Kaplan Meier survival curves in C57Bl6/J mice with PHx80% (n=73), mice with hepatocyte specific overexpression of HNF4α via AAV8-TBG-HNF4α (HNF4α-PHx80%; n=7) and mice with hepatocyte specific overexpression of HNF4α and G-CSF injections (HNF4α/G-CSF- PHx80%; n=8). Log-rank Mantel-Cox test. **(B)** Quantification of the PAS-stained area/total slide area on POD3 in GFP-PHx80% (n=7), HNF4α-PHx80% (n=4) and HNF4α/G-CSF-PHx80% (n=4). **(C)** Representative images of the regenerating livers on POD3 in GFP-PHx80%, HNF4α- PHx80% and HNF4α/G-CSF-PHx80%. In blue DAPI staining of the hepatocyte nuclei, in grey PAS hepatocyte glycogen storage, in green Ki67(+) hepatocytes and in orange pHH3(+) hepatocytes. Red arrow points to PAS^+^pHH3^+^ hepatocytes, white arrow points to PAS^-^pHH3^+^ hepatocytes. (Bar size = 20µm). **(D)** Quantification of Ki67^+^PAS^+^ hepatocytes on POD3 in GFP- PHx80% (n=6), HNF4α-PHx80% (n=3) and HNF4α/G-CSF-PHx80% (n=4). GFP-PHx80% vs HNF4α/G-CSF-PHx80% (\**p=0.03*) **(E)** Quantification of pHH3^+^PAS^+^ on POD3 in GFP-PHx80% (n=6), HNF4α-PHx80% (n=3) and HNF4α/G-CSF-PHx80% (n=4). GFP-PHx80% vs HNF4α- PHx80%; GFP-PHx80% vs HNF4α/G-CSF-PHx80% (\*\**p= 0.002* and \*\**p=0.001* respectively).

### Combination of G-CSF treatment and rescue of hepatocyte function increases survival in mice after SFSS-setting hepatectomy

HNF4α is a key transcription factor that regulates positively the epithelial phenotype and the expression of metabolic genes while it impacts negatively the proliferation in hepatocytes (*33*). As a high hepatocyte proliferative index was associated with poor function in SFSS-remnants, we injected AAV8-TBG-HNF4α virus to specifically overexpress HNF4α in hepatocytes to maintain terminal functional differentiation and prevent proliferation. We confirmed that HNF4α expression was upregulated and stable (Suppl Fig 3A) compared to vehicle controls (AAV8-TBG-GFP controls). HNF4α overexpression did not impact either hepatocyte proliferation (Suppl Fig 3B+C+D) or liver function in the SFSS-remnants (Suppl Fig 3E). However, double IF staining of PAS/pHH3 did show significantly more proliferating hepatocytes with conserved glycogen storage (reflecting hepatocyte function) (Suppl Fig 3F). This observation highlights that, in our model, HNF4α overexpression boosted function in the proliferating hepatocytes. However, this was not enough to sustain animal survival in the early phase of liver regeneration (Suppl Fig 3G).

In contrast, mice receiving the combined HNF4α/G-CSF treatment) showed a 3-day postoperative survival rate of 87.5% whereas survival of those injected with HNF4α alone was 57% (Fig 6A). Out of 7 surviving mice, we excluded 3 mice from further analysis due to high necrosis and microsteatosis on histology. As mentioned above, HNF4α overexpression, whether combined or not with G-CSF, did not alter hepatocyte proliferation, as assessed by Ki67 and pHH3 staining and mitotic counts (Suppl Fig 4A+B+C). However, the HNF4α/G-CSF-PHx80% group showed a high percentage of PAS staining compared to controls, although not significantly different (Fig 6B). We hypothesized that HNF4α overexpression forces hepatocyte function during the early phases of the cell cycle, thereby improving liver function during regeneration. We performed a double IF staining for Ki67 and pHH3 to identify hepatocytes in the early G1/S phase (Ki67^+^/pHH3^-^) or later G2/M (Ki67^+^/pHH3^+^) cell cycle stage, along with a PAS staining to assess their function (Fig 6C). The combined treatment resulted in a significantly higher number of PAS^+^ and Ki67^+^ “hybrid” hepatocytes compared to controls (p=0.03) (Fig 6D). Hepatocytes in late cell-cycle stages (pHH3^+^) exhibited increased metabolic activity with HNF4α overexpression compared to controls (GFP- PHx80% *vs* HNF4α-PHx80% or HNF4α/G-CSF-PHx80%, p=0.002 and p=0.001 respectively) (Fig 6E). The presence of such HNF4α-induced ‘hybrid’ hepatocytes combined to G-CSF treatment was associated with reduced plasma levels of total and indirect bilirubin (Fig 7A+B) while glycemia tended to be higher (Fig 7C). Finally, even though we observed no impact in albumin plasmatic levels in the treated animals (Fig 7D), expression of genes reflecting the overall hepatocyte function was slightly increased, with expression of *Ttr* being significant (Fig 7E). The liver function index, which takes all functional parameters assessed into account, was significantly higher with the combined treatment compared to controls (p=0.02) (Fig 7F).

**Fig. 7.**
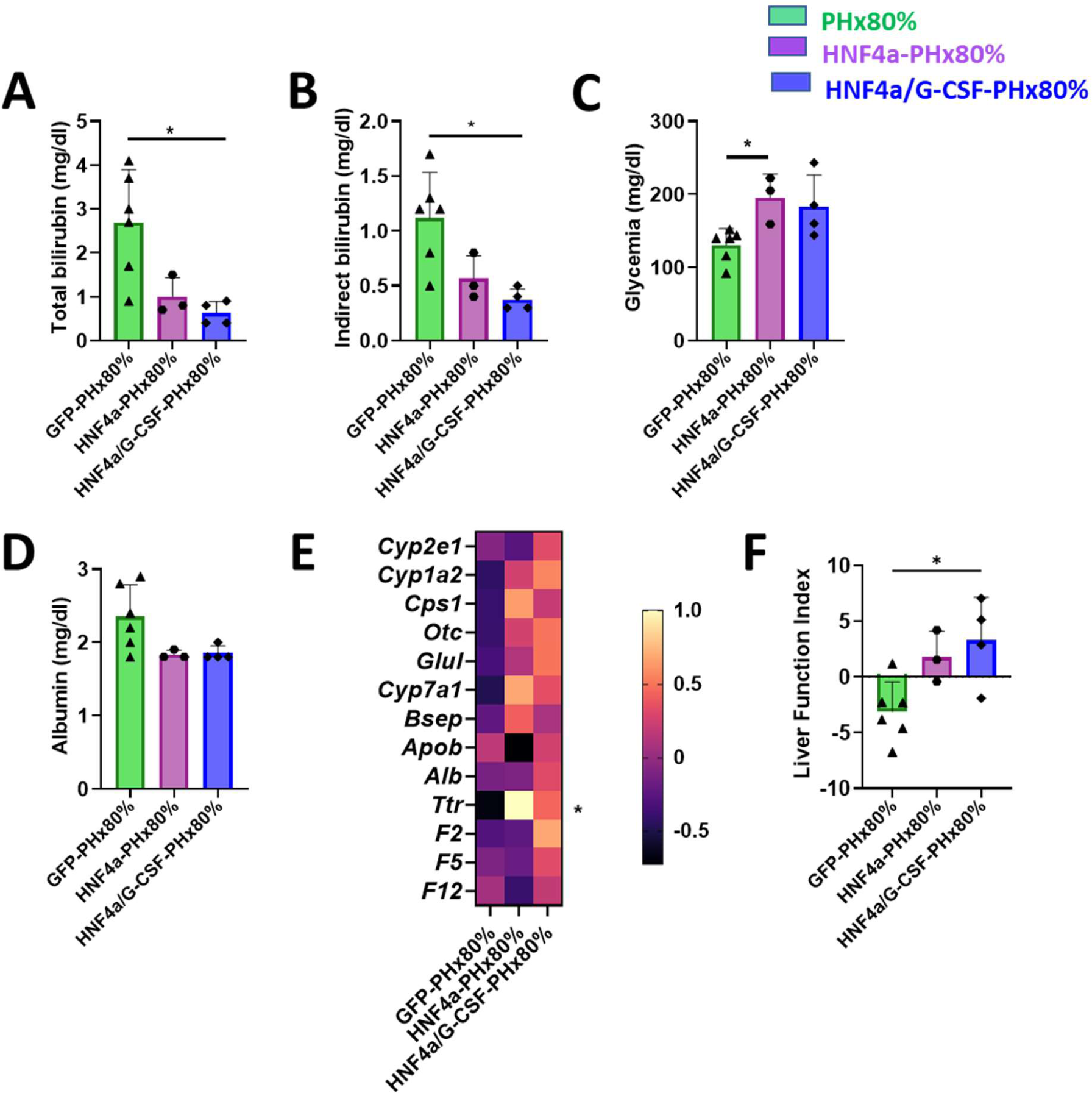
G-CSF treatment combined HNF4α overexpression increase function in mice after extended hepatectomy. **(A)** Total bilirubin plasma levels (mg/dl) on POD3 in GFP-PHx80% (n=6), HNF4α-PHx80% (n=3) and HNF4α/G-CSF-PHx80% (n=4). GFP-PHx80% vs HNF4α/G-CSF-PHx80% (\**p=0.01*); **(B)** indirect bilirubin plasma levels (mg/dl) (\**p=0.01*); **(C)** glucose plasma levels (mg/dl) ( (\**p=0.05*) and **(D)** albumin plasma levels (g/dl) in SFSS-setting hepatectomy on POD3 in GFP- PH80% (n=6), HNF4α-PHx80% (n=3) and HNF4α/G-CSF-PHx80% (n=4). **(E)** Heatmap of genes expressions reflecting hepatocyte function on POD3. Z-score was calculated from DeltaCT and plotted on the heatmap. GFP-PHx80% (n=7), HNF4α-PHx80% (n=3), HNF4α/G-CSF-PHx80% (n=4). **(F)** Liver function index (see methods for calculation) on POD3 in GFP-PHx80% (n=6), HNF4α-PHx80% (n=3) and HNF4α/G-CSF-PHx80% (n=4). GFP-PHx80% vs HNF4α/G-CSF- PHx80% (\**p=0.02*).

Thus, HNF4a treatment did not influence proliferation but stabilized the function of proliferating hepatocytes. The better survival in the HNF4α/G-CSF hepatectomized mice, together with the maintained function, suggests that both vascular remodeling and maintenance of hepatocyte function are key to prevent liver failure in the rapidly regenerating liver.

## DISCUSSION

Surgical resection is a crucial therapeutic tool for the treatment of primary or secondary liver cancers, providing the most favorable opportunity for achieving long-term survival (*1, 2*). However, despite improvements in surgical procedures and peri-operative care that expand the frontiers of resection or LT, hepatic surgery still carries some risk of post-operative liver failure, a condition associated with high morbidity and mortality (*9*). The larger the resection, the higher the risk of post-hepatectomy liver failure (*3*). Hence, the modulation of the future liver remnant before hepatectomy is currently the standard of care, a concept called ‘regenerative liver surgery’ (*34*). Yet, the clinical reality reveals that rapid volume recover is often linked to impaired liver function (*35*).

Portal hyperperfusion, arterial constriction with resulting hypoxia in the liver remnant, hepatocyte hyperproliferation and lobular disorganization, all contribute to postoperative liver failure (*14, 24*).

As such, vascular damage of the remnant emerges as leading cause of the SFSS. Yet, whether liver failure arises indirectly from microvascular injury and lobular disruption, as supported by observations of SFSS pathologic specimens in humans (*10, 19*), or directly from hepatocyte dysfunction remains unresolved. The present study provides evidence that hypoxia remodels the vasculature and mitigates hepatocyte proliferation to favor hepatocyte function after extended hepatectomy. It also shows that a two-pronged strategy, addressing simultaneously the preservation of function and the reduction of vascular harm in the regenerating organ, is essential to prevent mortality due to the small-for-size syndrome.

In a rodent model of major hepatectomy (PHx80%), post-hepatectomy mortality was high, and occurred mostly during the early phase of liver regeneration. In line with the literature (*14, 18*), we confirmed vascular damage indicated by sinusoidal collapse, intraparenchymal hemorrhage and infarcts, associated with poor liver function. As hypoxia is a master regulator of angiogenesis (*36–38*), we exposed mice to continuous hypoxia (FiO2 11%) immediately after extended hepatectomy for 3 consecutive days. Hypoxia post-extended hepatectomy was well tolerated, prevented vascular collapse, and increased microvascular density in the regenerating liver. It curtailed excessive shear stress-induced vessel denudation limiting parenchymal hemorrhage, while enhancing lobular perfusion and architecture. This paralleled with a high LSEC proliferation early after liver resection, which was not observed in control SFS-livers. This supports that the vascular damage in the SFS remnant results from a disequilibrium between the inevitable injury due to the increased shear stress and the delayed vascular remodeling.

It is accepted that endothelial progenitor cells (EPC) by integrating the existing vasculature and releasing growth factors (*39, 40*) are key for optimal liver regeneration (*32, 41, 42*). However, recruitment of EPC during liver regeneration is controversial. As nicely demonstrated by Singhal in 2018, endothelial fitness conditions the recruitment of endothelial progenitors (*28*). Indeed, EPCs integrated the vascular network when it was damaged by X-ray, but did not after 70% hepatectomy as the procedure caused no major vascular damage. In the SFSS model, we observed sinusoidal injuries implying that extended hepatectomy compromised endothelial fitness. Our data confirmed that non-liver native endothelial cells were then recruited and engrafted in the regenerating liver. Of note, native and non-native LSEC proliferated equally at day 2 post-surgery (data not shown), suggesting that the 2 populations of LSEC contribute to the healing and the expansion of the vascular network. The origin of endothelial progenitors has not been explored in our study. The literature supports that they originate from the bone marrow (*41, 43*) though a vascular, or even a sinusoidal, origin is not excluded. Activation of the VEGF/SDF-1α axis in mice exposed to hypoxia may dictate mobilization and recruitment of progenitors into the liver remnant, in line with previous reports (*29, 31*).

Endothelial progenitors also support regeneration by the release of hepatocyte growth factor (HGF), wingless-type MMTV integration site family member 2 (Wnt2) and angiopoietin-2 (*31, 44*), necessary for hepatocyte proliferation and sinusoidal reconstruction. The observation by Wang et al. in a 70% PHx model pointed out the essential role of bone-marrow-derived-EPC for regeneration (*29, 31*). Yet, in our study, hypoxia-induced recruitment of EPCs was associated with restrained hepatocyte proliferation but improved liver function in the hypoxic remnant. Similarly, increased mobilization and engraftment of bone marrow progenitors by G-CSF, although it triggered vascular renewal, was not associated with hepatocyte hyperproliferation. This indicates that the environment may influence the biology of recruited EPCs. It will therefore be of interest to further analyze the secretome and interactome of the recruited LSEC after a SFSS-setting hepatectomy, especially with exposure to hypoxia, to understand how they communicate with parenchymal liver cells to influence functional regeneration.

Liver resection induces a synchronized proliferation of hepatocytes (*45, 46*). The proportion of hepatocytes entering the cell cycle depends on the extent of liver resection (*47, 48*). Hence, extensive hepatectomy triggers a large proliferative response, with rapid mass recovery of the liver remnant. Using single cell RNA sequencing after a classical 70% partial hepatectomy, it was shown that hepatocytes segregate into proliferative and non-proliferative phenotypes. Replicative hepatocytes are less equipped for metabolic functions than the non-replicative ones, indicating a division of labor of hepatocytes during regeneration (*49–51*). Thus, rapid and high hepatocyte proliferation causes lack of function, as observed in our model of extended hepatectomy, as well as in human studies where rapid regeneration after major hepatectomy of LT of small grafts independently predicts the SFSS (*21, 23, 52*).

Hypoxia, besides impacting the liver microvasculature, restrained hepatocyte proliferation and favored function, while G-CSF, although it improved vascular remodeling, did not significantly mitigate hepatocyte proliferation and did not prevent liver failure. HNF4α is a direct repressor of cyclin D1 and a master transcription factor for hepatocyte identity (*53, 54*). In mice lacking HNF4α, hepatocyte proliferate spontaneously. After partial hepatectomy, despite a high proliferative index, the mice die, likely because of insufficient hepatocyte function, as reflected by elevated bilirubin (*55*). To test if a larger pool of quiescent and functionally active hepatocytes is needed to support functional recovery after a SFSS-setting hepatectomy, we overexpressed HNF4α in hepatocytes. In our hands, overexpression of HNF4α did not impact proliferation after 80% hepatectomy. However, we observed an increased number of metabolically active yet proliferating parenchymal cells. The increase in hepatocytes with a hybrid ‘proliferative and functional’ phenotype was paralleled by a slight improvement in liver function, although insufficiently to sustain animal survival. When combined to G-CSF treatment, HNF4α overexpression significantly improved functional liver recovery as well as the survival of the animal. Taken together, our data suggest that rapid functional regeneration may be achieved by blocking the entrance into the cell cycle as observed in hypoxic animals, or by supporting hepatocyte function during the cell cycle as imposed by overexpression of the hepatocyte master regulator HNF4α.

In conclusion, our research highlights the essential role of vascular remodeling in maintaining lobular architecture and perfusion to avoid the small-for-size syndrome. Nonetheless, our study accentuates that the restoration of function holds a pivotal position following extended hepatectomy. Achieving a metabolic and proliferative equilibrium in hepatocytes emerges as a critical factor for functional regeneration in small for size setting hepatectomy scenarios. This provides an explanation for the disparity between rapid volume recovery and functional outcomes in clinical studies. In addition, our data support that adopting a dual strategy that focuses on preserving function in proliferating hepatocytes while simultaneously addressing vascular damage holds the potential to prevent liver failure and contribute to the advancement of therapeutic approaches.

## MATERIALS AND METHODS

### Animals

All animal experiments were conducted in accordance with the European regulations and guidelines for humane care for laboratory animals. The study protocol was approved by the university ethics committee (2020/UCL/MD/020). Data of each individual animal were reported according to the ARRIVE guidelines. Animals were housed in a temperature- and humidity-controlled environment with a 12-h light/12-h dark cycle and ad libitum access to food and water.

All procedures on C57Bl/6J mice were carried out on male mice of minimum 8 weeks of age (>20 grams of body weight (BW)) and between 7 and 12am. For liver endothelial cell fate mapping, we used male and female Cdh5-PAC-Cre^ERT2^ mice crossed with the Rosa^26R^ mTomato/mGFP reporter mice (called from now on Cdh5-PAC-Cre^ERT2^XROSA^mT/mG^). To achieve *Cre-LoxP* recombination, mice received tamoxifen (T5648; Sigma, 100 mg/kg of body weight) intraperitoneally for 3 consecutive days (Suppl Fig 1). Surgery was performed not sooner than 2 weeks after tamoxifen injection.

### Surgical Procedures

During operation, animals were allowed to breathe spontaneously in a glass cylinder filled with a mixture of oxygen (2 l/min) and isoflurane (2.5%) (IsoFlo, Zoetis BelgiumSA, Louvain-la-Neuve, Belgium) for anesthesia. All animals had a median laparotomy after subcutaneous injection of 5cc of warm physiological sterile solution to increase the circulatory blood volume.

***70% Partial Hepatectomy (PHx70%):*** left lateral and median lobes were resected. As the procedure does not associate with substantial loss of liver function or mortality in rodents (*20, 56*), this model is used, when needed, as control for ‘physiological’ liver regeneration compared to extended partial hepatectomy.

***80% partial hepatectomy or extended, SFSS-setting, hepatectomy (PHx80%):*** We established an extended partial hepatectomy model, in which all but the superior right lobe (SRL∼12,65% of the total liver volume) and the caudate lobe (CL∼6% of the total liver volume) were removed, using microscope-assisted dissection (20x magnification) as previously described (*57–59*). As such, the liver remnant (LR) accounts for around 20 % of the total liver volume.

***Exposure to hypoxia after 80% partial hepatectomy (PHx80%-HC):*** Immediately after surgery, animals were placed into hypoxic chambers and exposed to continuous hypoxia (Fi02 of 11%) for 3 consecutive days (i.e., during the early phase of liver regeneration). Previous studies, mainly in lung and heart disease animal models, have demonstrated that the reduction of the inspired oxygen fraction to 11% (FiO2) is well tolerated and activates hypoxia inducible factors (HIF) and neoangiogenesis (*60, 61*). Control animals inhaled the surrounding air, with inspired oxygen fraction of 21% (PHx80%).

***Treatment with Granulocyte Colony Stimulating Factor (G-CSF-PHx80%):*** Mice received 3 sub-cutaneous injections of G-CSF (250µg/kg of body weight): the first two ones the day before the operation at 10 am and 6 pm, the third one at completion of the extended liver resection.

***HNF4***α ***virus-mediated hepatocyte overexpression (HNF4***α***-PHx80%):*** We used intravenous injection of AAV8-TBG-HNF4α virus (RefSeq: NM_008261) from Vector Biolabs (AAV- 261497, Malvern, USA) to overexpress HNF4α specifically in hepatocytes. We injected 2.5*10^11^ GC of AAV8-TBG-HNF4α in PBS in our experimental groups or AAV8-TBG-eGFP in controls, 4 days before 80% liver resection.

In the figure legends, we report the number of animals or samples on which each analysis was performed.

### In vivo liver perfusion with fixable fluorescein-Dextran

To evaluate liver sinusoidal perfusion in vivo, fixable fluorescein-dextran (D1820, Invitrogen) (0.03mg/g of body weight) was injected into the inferior vena cava in anaesthetized mice 3 days after hepatectomy. After 10 minutes, the liver was harvested, fixed in a 4% buffered formaldehyde solution, embedded in 6% gelatin, and then cut with a vibratome. 200µm-thick slices were analyzed with a fluorescent confocal microscope (LSM980-MP Airyscan, Zeiss, Germany). Non-perfused area was defined as an area where fluorescein-dextran was not observed and vascular leak as an area where fluorescein-dextran had diffused outside the sinusoidal bed. Analysis and measurements were done on anonymized samples using ImageJ software (National Institutes of Health, Bethesda, Maryland, USA). Results are all expressed as the mean of measurements of minimum 5 images per mouse.

### Plasma analysis

Plasma was collected after centrifugation of whole-blood samples at 3,000g for 10minutes at 4°C. We measured albumin, total and indirect bilirubin and glucose concentrations by colorimetry using a commercially available kit (Fujifilm) and an automatic biochemical analyzer (DRI-CHEM NX500 system®). U-Plex assay (K15069L, Meso Scale Diagnostic, Rockville, Maryland, USA) was chosen to measure serum concentration of Granulocyte-macrophage colony-stimulating factor (GM-CSF), stromal cell-derived factor-1α (SDF-1α) and vascular endothelial growth factor A (VEGF-A). Factor V serum protein concentration was assessed by ELISA following manufacturer’s instructions (Mouse factor V ELISA kit orb409284, Biorbyt).

### Histologic analysis

We performed fluorescent immunohistochemistry on 4 µm-thick liver slices. We used antiCD31 antibody (77699S, Cell Signaling, diluted 1/200) to detect endothelial cells. We manually measured the diameter of the sinusoids as the smallest diameter of the transversally cut vessels (Feret’s diameter) as previously described (*62*). We draw a line horizontally across the image, avoiding large vessels, and counted the number of sinusoids crossing the line as a proxy for sinusoidal density. Analysis and measurements were done on anonymized samples using ImageJ software (National Institutes of Health, Bethesda, Maryland, USA). Results are all expressed as the mean of measurements on minimum 5 images per mouse.

CD31 immunofluorescent staining combined to Ki67 (12202S, Cell Signaling, diluted 1/100) was used to identify endothelial cells engaged in the cell cycle. Endothelial cell proliferation was expressed as percentage of CD31^+^ cells expressing Ki67 in relation to the entire population of endothelial cells (endothelial cell proliferative index). We used the VisioPharm software (VisioPharm, Hørsholm, Denmark) on 20X magnification tissue scans.

We used Cdh5-PAC-Cre^ERT2^XRosa^mT/mG^ mice to label LSEC. With fluorescent microscopy, we identified VE-Cadherin by immunofluorescence (AF2002, dilution 1/100, RND Systems) along with the endogenous GFP signal. We measured the proportion of VE-Cadherin^+^ cells expressing GFP to assess the labeling efficiency of the system (and further confirmed it by flow cytometry) and determined the involvement of endothelial progenitors or cell recruitment to the vascular remodeling during regeneration.

To evaluate hepatocyte proliferation, we counted mitotic hepatocytes on hematoxylin/eosin-stained liver slides (mean of minimum 5 random 20X magnification fields per remnant liver). We also used immunofluorescence with Hepatocyte Nuclear Factor 4a (HNF4α) (1/350, PP-H1415- 00, Perseus Proteomics) to define hepatocytes in combination with phospho-histone H3 (pHH3) (9701S, Cell Signaling, diluted 1/200) or Ki67 (12202S, Cell Signaling, diluted 1/100). DAPI (D9542, Sigma) counterstained all nuclei.

Periodic acid-Schiff (PAS) (periodic acid, Merck-Millipore, 100524002, Schiff’s reagent, Sigma-Aldrich, 3952016) staining was used to identify glycogen storage within hepatocytes. PAS positive area was quantified using an artificial intelligence-trained classifier on HALO software and expressed as a percentage of the total tissue area. To assess PAS staining in proliferating hepatocytes, we performed a triple staining: Ki67 and pHH3 using TSA dyes; then, PAS staining was performed as mentioned above. The fluorescent signal of PAS was acquired in the 555nm channel.

To define hepatocyte islets, E-Cadherin (ab76319, Abcam, diluted 1/200), a marker of hepatocyte basolateral membrane, and CD31 (77699S, Cell Signaling, diluted 1/100) were used for the staining. We captured 10 images (at 20X magnification) around the central or portal tract and counted the hepatocytes or groups of hepatocytes that had no contact with any sinusoid.

### Gene expression analyses

We quantified the gene expression of proteins associated with hepatocyte overall function by qPCR on RNA extracted from the entire liver remnant. Total RNA was extracted from frozen liver samples by using TRIZOL Isolation Reagent (15596018, ThermoFischer). cDNA was synthesized from 1 µg of RNA. Real-time PCR gene expression analysis was performed using Rotor gene Q device and software (Qiagen) using SYBRGreen (4309155, Applied Biosystems) and gene specific primer pairs for transcripts of interest (Suppl. Table 1). Ribosomal protein L19 mRNA was chosen as an invariant standard. All experimental samples were run in duplicate. Z-score was calculated from the DeltaCT for all genes and expressed on a heatmap.

### Liver function index

Z-scores were calculated for each functional parameter assessed (total, direct, indirect bilirubin, PAS^+^ area, circulating albumin, functional genes heatmap, glycemia) to standardize the datasets.

The liver function index was obtained by summing the parameters positively correlated with function and subtracting the negatively correlated parameters.

### Statistical analysis

GraphPad Prism software (San Diego, CA, USA) was used to do the graphics and the statistics. In graphs, we report individual data (dots), the mean ± standard error (SEM). Survival curves were analyzed using the LogRang test (Mantel-Cox). Unpaired two-tailed t-test was used for simple comparison or one-way and two-way ANOVA followed by Bonferroni’s post-hoc correction for multiple comparison. Statistical significance was assumed for p values <0.05 (*p<0.05; **p<0.01; ***p<0.001; **** p<0.0001).

## List of Supplementary Materials

Fig S1-S4 (see supplementary figures section). Table S1 (See supplementary table section)

## Acknowledgments

We thank Dr Caroline Bouzin and the I2P imaging platform (IREC, UCL, Brussels, Belgium) for IHC and morphometrical quantification, Patrick Vandersmissen for Dextran perfusion experiment (IREC, UCL, Brussels, Belgium), Natacha Feza-Bingui and Boris Pirlot (GAEN Lab, IREC, UCL, Brussels, Belgium) for animal breeding, experiments and care and technical support. We also thank Prof Anne Sonet (haematology, CHU-UCL Namur, site of Godinne) for advice on GCSF administration and Prof. Yves Horsmans (St Luc university Hospital, UCL, Brussels, Belgium) for critical review of the manuscript.

## Funding

AD is a postdoctoral fellow from the Fond national de la recherche scientifique (FNRS), Belgium; MDR is a FNRS PhD fellow;

RM is a FNRS postdoctoral fellow;

The study was supported by research grants from the Foundation Mont Godinne (n°2022-BR-02 to AD);

And from the FNRS (CDR grant n°40014061 to AD and CDR grant n°J.0130.20 ‘Hypoxia for optimal regeneration’ to IL).

## Author contributions

Conceptualization: AD, MDR, IL Investigation: AD, MDR, AF, RM, LC, CB, IL Visualization: AD, MDR, IL

Funding acquisition: AD, IL

Writing – original draft: MDR, AD, IL

Writing – review & editing: AD, MDR, RM, IL

## Data and materials availability

All data are available in the main text or the supplementary materials.

## References and Notes

1. R. T. Poon, S. T. Fan, C. M. Lo, C. L. Liu, C. M. Lam, W. K. Yuen, C. Yeung, J. Wong, D. M. Nagorney, J. M. Henderson, H. A. Pitt, R. T. Poon, Improving perioperative outcome expands the role of hepatectomy in management of benign and malignant hepatobiliary diseases: Analysis of 1222 consecutive patients from a prospective database. Ann. Surg. 240, 698–710 (2004).

2. H. Imamura, Y. Seyama, N. Kokudo, T. Aoki, K. Sano, M. Minagawa, Y. Sugawara, M. Makuuchi, Single and multiple resections of multiple hepatic metastases of colorectal origin. Surgery 135, 508–517 (2004).

3. W. R. Jarnagin, M. Gonen, Y. Fong, R. P. DeMatteo, L. Ben-Porat, S. Little, C. Corvera, S. Weber, L. H. Blumgart, Improvement in perioperative outcome after hepatic resection: analysis of 1,803 consecutive cases over the past decade. Ann. Surg. 236, 397–406; discussion 406-7 (2002).

4. P.-A. Clavien, H. Petrowsky, M. L. DeOliveira, R. Graf, Strategies for Safer Liver Surgery and Partial Liver Transplantation. N. Engl. J. Med. 356, 1545–1559 (2007).

5. A. Amer, C. H. Wilson, D. M. Manas, Liver transplantation for unresectable malignancies: Beyond hepatocellular carcinoma. Eur. J. Surg. Oncol. 45, 2268–2278 (2019).

6. A. T. W. Song, V. I. Avelino-Silva, R. A. A. Pecora, V. Pugliese, L. A. C. D’Albuquerque, E. Abdala, Liver transplantation: Fifty years of experience. World J. Gastroenterol. 20, 5363–5374 (2014).

7. K. Fukazawa, Y. Yamada, S. Nishida, T. Hibi, K. L. Arheart, E. A. Pretto, Determination of the safe range of graft size mismatch using body surface area index in deceased liver transplantation. Transpl. Int. 26, 724–733 (2013).

8. F. Dahm, P. Georgiev, P. A., et al Clavien, Small-for-size syndrome after partial liver transplantation: Definition, mechanisms of disease and clinical implications. Am. J. Transplant. 5, 2605–2610 (2005).

9. N. N. Rahbari, O. J. Garden, R. Padbury, M. Brooke-Smith, M. Crawford, R. Adam, M. Koch, M. Makuuchi, R. P. Dematteo, C. Christophi, S. Banting, V. Usatoff, M. Nagino, G. Maddern, T. J. Hugh, J.-N. Vauthey, P. Greig, M. Rees, Y. Yokoyama, S. T. Fan, Y. Nimura, J. Figueras, L. Capussotti, M. W. Büchler, J. Weitz, Posthepatectomy liver failure: A definition and grading by the International Study Group of Liver Surgery (ISGLS). Surgery 149, 713–724 (2011).

10. M. Golriz, A. Majlesara, S. El Sakka, M. Ashrafi, J. Arwin, N. Fard, H. Raisi, A. Edalatpour, A. Mehrabi, Small for Size and Flow (SFSF) syndrome: An alternative description for posthepatectomy liver failure. Clin. Res. Hepatol. Gastroenterol. 40, 267–275 (2016).

11. N. N. Rahbari, O. J. Garden, R. Padbury, G. Maddern, M. Koch, T. J. Hugh, S. T. Fan, Y. Nimura, J. Figueras, J.-N. N. Vauthey, M. Rees, R. Adam, R. P. DeMatteo, P. Greig, V. Usatoff, S. Banting, M. Nagino, L. Capussotti, Y. Yokoyama, M. Brooke-Smith, M. Crawford, C. Christophi, M. Makuuchi, M. W. Büchler, J. Weitz, Post-hepatectomy haemorrhage: A definition and grading by the International Study Group of Liver Surgery (ISGLS). Hpb 13, 528–535 (2011).

12. V. Smyrniotis, G. Kostopanagiotou, A. Kondi, E. Gamaletsos, K. Theodoraki, D. Kehagias, K. Mystakidou, J. Contis, Hemodynamic interaction between portal vien and hepatic artery flow in small-for-size split liver transplantation. Transpl. Int. 15, 355–360 (2002).

13. M. Sainz-Barriga, L. Scudeller, M. G. Costa, B. de Hemptinne, R. I. Troisi, Lack of a correlation between portal vein flow and pressure: Toward a shared interpretation of hemodynamic stress governing inflow modulation in liver transplantation. Liver Transplant. 17, 836–848 (2011).

14. A. J. Demetris, D. M. Kelly, B. Eghtesad, P. Fontes, J. Wallis Marsh, K. Tom, H. P. Tan, T. Shaw-Stiffel, L. Boig, P. Novelli, R. Planinsic, J. J. Fung, A. Marcos, Pathophysiologic observations and histopathologic recognition of the portal hyperperfusion or small-for-size syndrome. Am. J. Surg. Pathol. 30, 986–993 (2006).

15. C. Eipel, K. Abshagen, B. Vollmar, Regulation of hepatic blood flow: The hepatic arterial buffer response revisited. World J. Gastroenterol. 16, 6046–6057 (2010).

16. G. K. Michalopoulos, Liver regeneration. J. Cell. Physiol. 213, 286–300 (2007).

17. P. F. Wang, C. H. Li, Y. W. Chen, A. Q. Zhang, S. W. Cai, J. H. Dong, Preserving hepatic artery flow during portal triad blood inflow occlusion improves remnant liver regeneration in rats after partial hepatectomy. J. Surg. Res. 181, 329–336 (2013).

18. D. M. Kelly, A. J. Demetris, J. J. Fung, A. Marcos, Y. Zhu, V. Subbotin, L. Yin, E. Totsuka, T. Ishii, M. C. Lee, J. Gutierrez, G. Costa, R. Venkataraman, J. R. Madariaga, Porcine partial liver transplantation: A novel model of the ?small-for-size? liver graft. Liver Transplant. 10, 253–263 (2004).

19. J. M. Asencio, J. Vaquero, L. Olmedilla, J. L. García Sabrido, “ Small-for-flow” syndrome: Shifting the “ size” paradigm. Med. Hypotheses 80, 573–577 (2013).

20. L. Yang, Y. Luo, L. Ma, H. Wang, W. Ling, J. Li, X. Qi, Q. Lu, K. Chen, Establishment of a novel rat model of different degrees of portal vein stenosis following 70% partial hepatectomy. Exp. Anim. 65, 165–173 (2016).

21. S. Gruttadauria, D. Pagano, R. Liotta, A. Tropea, F. Tuzzolino, G. Marrone, G. Mamone, J. W. Marsh, R. Miraglia, A. Luca, G. Vizzini, B. G. Gridelli, Liver Volume Restoration and Hepatic Microarchitecture in Small-for-Size Syndrome. Ann. Transplant. 20, 381–9 (2015).

22. M. Ninomiya, K. Shirabe, T. Terashi, H. Ijichi, Y. Yonemura, N. Harada, Y. Soejima, A. Taketomi, M. Shimada, Y. Maehara, Deceleration of Regenerative Response Improves the Outcome of Rat with Massive Hepatectomy. Am. J. Transplant. 10, 1580–1587 (2010).

23. J. Belghiti, G. Liddo, V. Raut, M. Zappa, S. Dokmak, V. Vilgrain, F. Durand, F. Dondéro, “Inherent limitations” in donors: Control matched study of consequences following a right hepatectomy for living donation and benign liver lesions. Ann. Surg. 255, 528–533 (2012).

24. M. De Rudder, A. Dili, P. Stärkel, I. A. Leclercq, Review critical role of lsec in post- hepatectomy liver regeneration and failure. Int. J. Mol. Sci. 22 (2021), doi:10.3390/ijms22158053.

25. A. Dili, C. Bertrand, V. Lebrun, B. Pirlot, I. A. Leclercq, Hypoxia protects the liver from Small For Size Syndrome: A lesson learned from the associated liver partition and portal vein ligation for staged hepatectomy (ALPPS) procedure in rats. Am. J. Transplant. 19, 2979–2990 (2019).

26. Ben Z. Stanger, Cellular Homeostasis and Repair in the Mammalian Liver Ben. Annu. Rev. Physiol. 176, 139–148 (2015).

27. Z. J. Liu, O. C. Velazquez, Hyperoxia, endothelial progenitor cell mobilization, and diabetic wound healing. Antioxidants Redox Signal. 10, 1869–1882 (2008).

28. M. Singhal, X. Liu, D. Inverso, K. Jiang, J. Dai, H. He, S. Bartels, W. Li, A. A. Abdul Pari, N. Gengenbacher, E. Besemfelder, L. Hui, H. G. Augustin, J. Hu, A. A. A. Pari, N. Gengenbacher, E. Besemfelder, L. Hui, H. G. Augustin, J. Hu, Endothelial cell fitness dictates the source of regenerating liver vasculature. J. Exp. Med. 215, 2497–2508 (2018).

29. L. Wang, X. Wang, L. Wang, J. D. Chiu, G. van de Ven, W. A. Gaarde, L. D. DeLeve, Hepatic Vascular Endothelial Growth Factor Regulates Recruitment of Rat Liver Sinusoidal Endothelial Cell Progenitor Cells. Gastroenterology 143, 1555–1563.e2 (2012).

30. H. Chopra, M. K. Hung, D. L. Kwong, C. F. Zhang, E. H. N. Pow, Insights into endothelial progenitor cells: Origin, classification, potentials, and prospects. Stem Cells Int. 2018 (2018), doi:10.1155/2018/9847015.

31. L. D. Deleve, X. Wang, L. Wang, VEGF-sdf1 recruitment of CXCR7+ bone marrow progenitors of liver sinusoidal endothelial cells promotes rat liver regeneration. Am. J. Physiol. - Gastrointest. Liver Physiol. 310, G739–G746 (2016).

32. R. Harb, G. Xie, C. Lutzko, Y. Guo, X. Wang, C. K. Hill, G. C. Kanel, L. D. DeLeve, Bone Marrow Progenitor Cells Repair Rat Hepatic Sinusoidal Endothelial Cells After Liver Injury. Gastroenterology 137, 704–712 (2009).

33. C. Walesky, U. Apte, Role of hepatocyte nuclear factor 4α (HNF4α) in cell proliferation and cancer. Gene Expr. 16, 101–108 (2015).

34. J. Heil, M. Schiesser, E. Schadde, Current trends in regenerative liver surgery: Novel clinical strategies and experimental approaches. Front. Surg. 9, 1–14 (2022).

35. P. B. Olthof, E. Schadde, K. P. van Lienden, M. Heger, K. de Bruin, J. Verheij, R. J. Bennink, T. M. van Gulik, Hepatic parenchymal transection increases liver volume but not function after portal vein embolization in rabbits. Surg. (United States) 162, 732–741 (2017).

36. B. L. Krock, N. Skuli, M. C. Simon, Hypoxia-Induced Angiogenesis: Good and Evil. Genes and Cancer 2, 1117–1133 (2011).

37. M. Schäfer, N. Ewald, C. Schäfer, A. Stapler, H. M. Piper, T. Noll, Signaling of hypoxia-induced autonomous proliferation of endothelial cells. FASEB J. 17, 1–23 (2003).

38. Z. Qing, H. Huang, Q. Luo, J. Lin, S. Yang, T. Liu, Z. Zeng, T. Ming, Hypoxia promotes the proliferation of mouse liver sinusoidal endothelial cells: miRNA-mRNA expression analysis. Bioengineered 12, 8666–8678 (2021).

39. X. jun Zhang, V. Olsavszky, Y. Yin, B. Wang, T. Engleitner, R. Öllinger, K. Schledzewski, P. S. Koch, R. Rad, R. M. Schmid, H. Friess, S. Goerdt, N. Hüser, C. Géraud, G. von Figura, D. Hartmann, Angiocrine Hepatocyte Growth Factor Signaling Controls Physiological Organ and Body Size and Dynamic Hepatocyte Proliferation to Prevent Liver Damage during Regeneration. Am. J. Pathol. 190, 358–371 (2020).

40. M. Winkler, T. Staniczek, S. W. Kürschner, C. D. Schmid, H. Schönhaber, J. Cordero, L. Kessler, A. Mathes, C. Sticht, M. Neßling, A. Uvarovskii, S. Anders, X. jun Zhang, G. von Figura, D. Hartmann, C. Mogler, G. Dobreva, K. Schledzewski, C. Géraud, P. S. Koch, S. Goerdt, Endothelial GATA4 controls liver fibrosis and regeneration by preventing a pathogenic switch in angiocrine signaling. J. Hepatol. 74, 380–393 (2021).

41. H. Fujii, T. Hirose, S. Oe, K. Yasuchika, H. Azuma, T. Fujikawa, M. Nagao, Y. Yamaoka, Contribution of bone marrow cells to liver regeneration after partial hepatectomy in mice. J. Hepatol. 36, 653–659 (2002).

42. L. Wang, X. Wang, G. Xie, L. Wang, C. K. Hill, L. D. Deleve, Liver sinusoidal endothelial cell progenitor cells promote liver regeneration in rats. J. Clin. Invest. 122, 1567–1573 (2012).

43. L. D. DeLeve, Liver sinusoidal endothelial cells and liver regeneration*J*. Clin. Invest. 123, 1861–1866 (2013).

44. B.-S. Ding, D. J. Nolan, J. M. Butler, D. James, A. O. Babazadeh, Z. Rosenwaks, V. Mittal, H. Kobayashi, K. Shido, D. Lyden, T. N. Sato, S. Y. Rabbany, S. Rafii, Inductive angiocrine signals from sinusoidal endothelium are required for liver regeneration. Nature 468, 310–315 (2010).

45. J. I. Fabrikant, The kinetics of cellular proliferation in regenerating liver. J. Cell Biol. 36, 551–565 (1968).

46. Y. Zou, Q. Bao, S. Kumar, M. Hu, G. Y. Wang, G. Dai, Four waves of hepatocyte proliferation linked with three waves of hepatic fat accumulation during partial hepatectomy-induced liver regeneration. PLoS One 7 (2012), doi:10.1371/journal.pone.0030675.

47. T. Kobayashi, H. Imamura, T. Aoki, Y. Sugawara, N. Kokudo, M. Makuuchi, Morphological Regeneration and Hepatic Functional Mass after Right Hemihepatectomy. Dig. Surg. 23, 44–50 (2006).

48. M. Meier, K. J. Andersen, A. R. Knudsen, J. R. Nyengaard, S. Hamilton-Dutoit, F. V. Mortensen, Liver regeneration is dependent on the extent of hepatectomy. J. Surg. Res. 205, 76– 84 (2016).

49. T. Chen, S. Oh, S. Gregory, X. Shen, A. M. Diehl, Single-cell omics analysis reveals functional diversification of hepatocytes during liver regeneration. JCI Insight 5 (2020), doi:10.1172/jci.insight.141024.

50. U. V. Chembazhi, S. Bangru, M. Hernaez, A. Kalsotra, Cellular plasticity balances the metabolic and proliferation dynamics of a regenerating liver. Genome Res. 31, 576–591 (2021).

51. S. Minocha, D. Villeneuve, L. Rib, C. Moret, N. Guex, W. Herr, Segregated hepatocyte proliferation and metabolic states within the regenerating mouse liver. Hepatol. Commun. 1, 871–885 (2017).

52. F. R. Pruvot, S. Truant, Major hepatic resection: from volumetry to liver scintigraphy. Hpb 18, 707–708 (2016).

53. H. Wu, T. Reizel, Y. J. Wang, J. L. Lapiro, B. T. Kren, J. Schug, S. Rao, A. Morgan, A. Herman, L. L. Shekels, M. S. Rassette, A. N. Lane, T. Cassel, T. W. M. Fan, J. C. Manivel, S. Gunewardena, U. Apte, P. Sicinski, K. H. Kaestner, J. H. Albrecht, A negative reciprocal regulatory axis between cyclin D1 and HNF4α modulates cell cycle progression and metabolism in the liver. Proc. Natl. Acad. Sci. U. S. A. 117, 17177–17186 (2020).

54. C. Cicchini, L. Amicone, T. Alonzi, A. Marchetti, C. Mancone, M. Tripodi, Molecular mechanisms controlling the phenotype and the EMT/MET dynamics of hepatocyte. Liver Int. 35, 302 (2015).

55. I. Huck, S. Gunewardena, R. Espanol-Suner, H. Willenbring, U. Apte, Hepatocyte Nuclear Factor 4 alpha (HNF4α) Activation is Essential for Termination of Liver Regeneration. Hepatology 70, 666 (2019).

56. C. Mitchell, H. Willenbring, A reproducible and well-tolerated method for 2/3 partial hepatectomy in mice. Nat. Protoc. 3, 1167–1170 (2008).

57. M. A. Aller, N. Arias, I. Prieto, S. Agudo, C. Gilsanz, L. Lorente, J. L. J. Arias, J. L. J. Arias, A half century (1961-2011) of applying microsurgery to experimental liver research. World J. Hepatol. 4, 199–208 (2012).

58. N. Madrahimov, O. Dirsch, C. Broelsch, U. Dahmen, Marginal Hepatectomy in the Rat. Ann. Surg. 244, 89–98 (2006).

59. P. N. Martins, Paulo Ney Aguiar, Hepatic lobectomy and segmentectomy models using microsurgical techniques. Microsurgery 28, 187–191 (2008).

60. M. K. Ball, G. B. Waypa, P. T. Mungai, J. M. Nielsen, L. Czech, V. J. Dudley, L. Beussink, R. W. Dettman, S. K. Berkelhamer, R. H. Steinhorn, S. J. Shah, P. T. Schumacker, Regulation of hypoxia-induced pulmonary hypertension by vascular smooth muscle hypoxia-inducible factor- 1α. Am. J. Respir. Crit. Care Med. 189, 314–324 (2014).

61. Y. Nakada, Z. Lu, D. C. Canseco, Z. Hu, C. C. Zhang, L. I. Szweda, G. Schiattarella, M. T. Kinter, S. Thet, H. Zhang, S. Abdisalaam, J. A. Hill, A. M. Shah, C. X. Santos, A. Asaithamby, C. Xing, O. Oz, H. A. Sadek, J. E. Faber, W. Kimura, R. J. Deberardinis, C. X. Santos, A. M. Shah, H. Zhang, J. E. Faber, M. T. Kinter, L. I. Szweda, C. Xing, Z. Hu, R. J. Deberardinis, G. Schiattarella, J. A. Hill, O. Oz, Z. Lu, C. C. Zhang, W. Kimura, H. A. Sadek, Hypoxia induces heart regeneration in adult mice. Nature 541, 222–227 (2016).

62. A. Dili, M. De Rudder, B. Pirlot, C. Bertrand, L. Dewachter, C. Bouzin, I. Leclercq, Hypoxia induced Angiogenesis Rescues Survival from Small for Size Syndrome (SFSS). Hpb 23, S103 (2021).

