## Supplementary material for "Vascular damage and excessive proliferation compromise liver function after extended hepatectomy in mice": Suppl Fig

**Supplementary figures**

### Suppl. Fig. 1

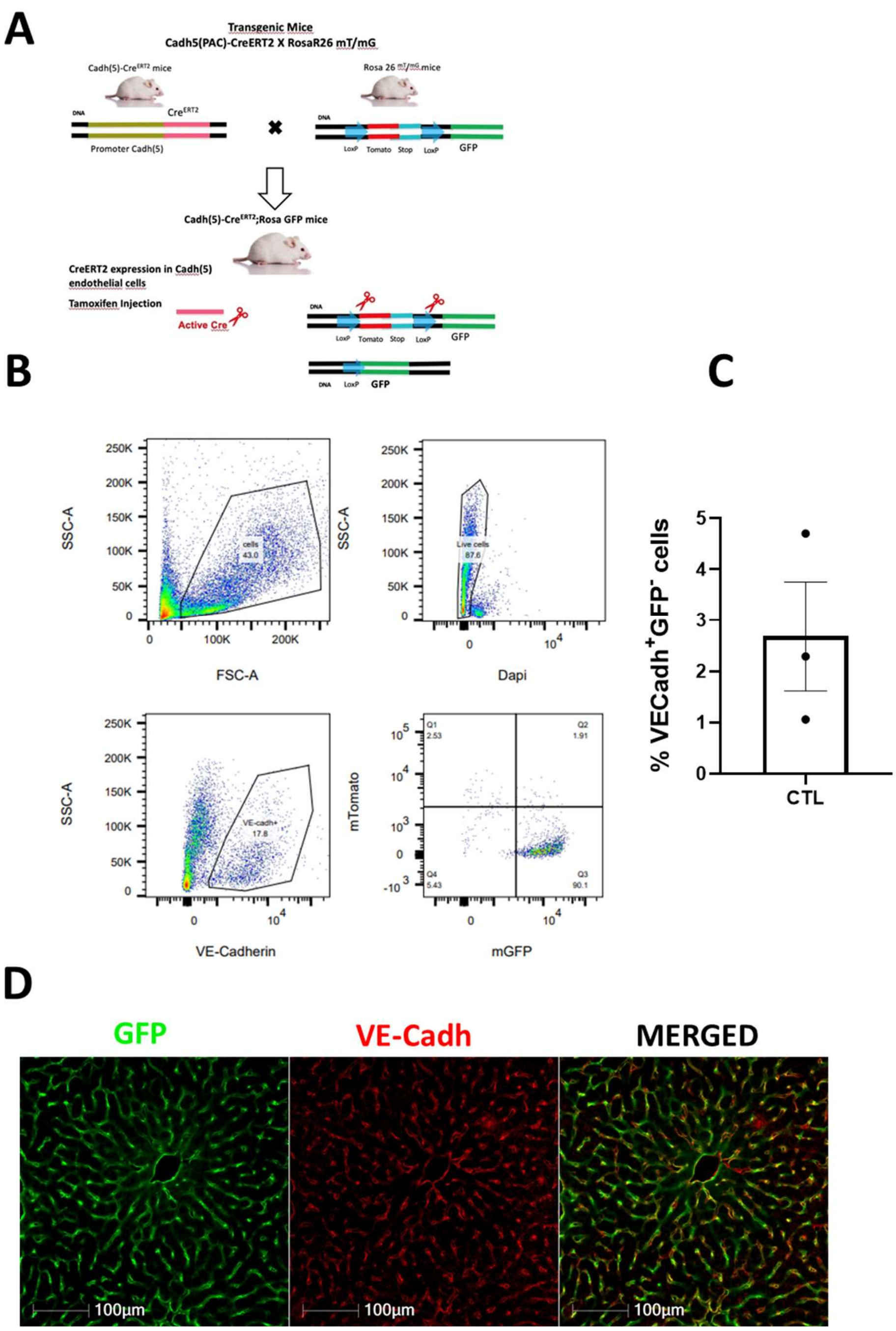

**Suppl. Fig. 1 Tamoxifen induces a high rate of recombination in liver endothelial cells in Cdh5-PAC-Cre$^{ERT2}$XROSA$^{mT/mG}$ mice.**

**(A)** Schematic representation of the Cdh5-PAC-Cre$^{ERT2}$xROSA$^{mT/mG}$ treated with Tamoxifen mice for liver endothelial cell fate mapping, Cdh5-PAC-Cre$^{ERT2}$xROSA$^{mT/mG}$ treated with Tamoxifen mouse model was used, undergoing the same study design as for C57Bl6/J mice. Briefly, before tamoxifen injection, all endothelial cells express mTomato. When tamoxifen is injected, endothelial cells expressing VECadherin (*Cdh5*) express the Cre recombinase. The active Cre enters the nucleus and excise the mTomato out of the DNA. After that, only endothelial cells (at the time of tamoxifen injection) express the mGFP. **(B)** Flow cytometry assessing mGFP and mTomato expression in VECadh$^{+}$ cells in Cdh5-PAC-Cre$^{ERT2}$XROSA$^{mT/mG}$ mice **(C)** Quantification of the percentage of GFP$^{-}$VECadh$^{+}$ cells (not recombined endothelial cells) in tamoxifen induced Cdh5-PAC-Cre$^{ERT2}$XROSA$^{mT/mG}$ mice (n=3). **(D)** Representative immunofluorescence image of the liver of tamoxifen induced Cdh5-PAC-Cre$^{ERT2}$XROSA$^{mT/mG}$ mice showing that nearly all endothelial cells (VECadh$^{+}$) are labelled by GFP$^{+}$ (Bar size = 100μm).

### Suppl. Fig. 2

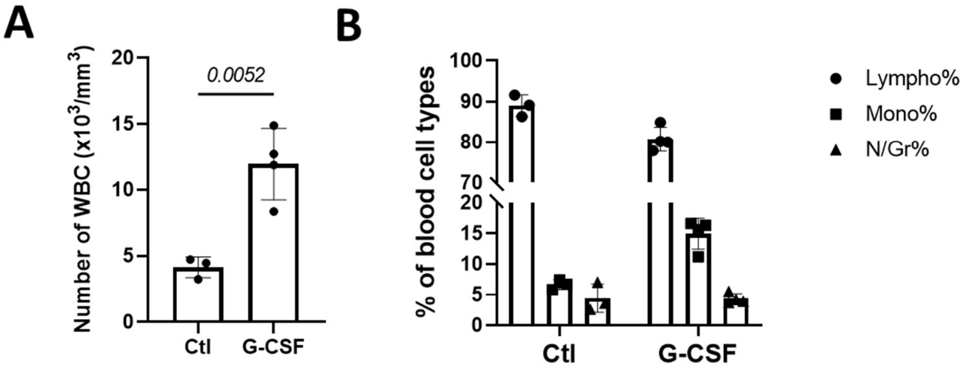


**Suppl. Fig. 2 G-CSF increases blood monocytes without disturbing the proportion of other blood cell types.**

**(A)** Quantification of the number of white blood cells in the blood of controls (n=3) and G-CSF (sacrificed one day after the last injection) treated mice (n=4). **(B)** Distribution (in percentage) of lymphocytes, monocytes and neutrophils/granulocytes in the blood of controls (n=3) and G-CSF treated mice (n=4).

#### Suppl. Fig. 3

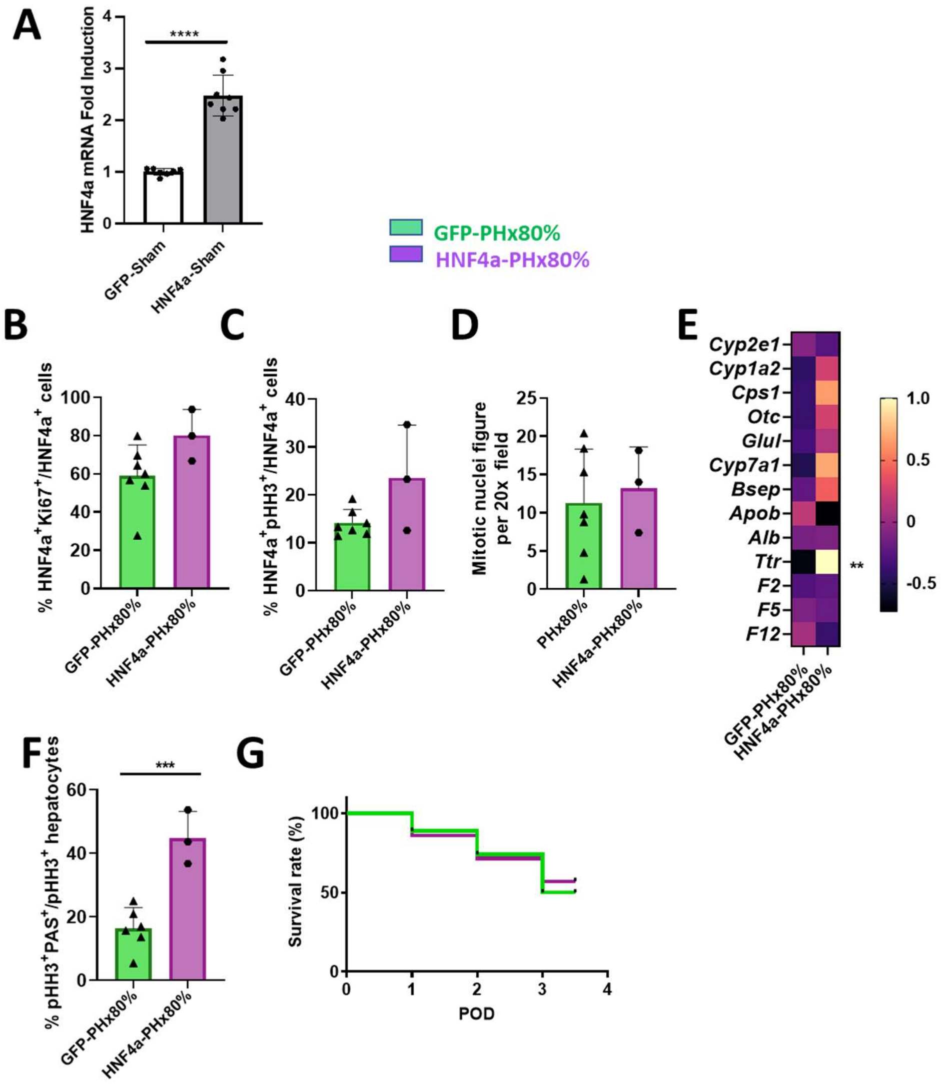
A
HNF4a mRNA Fold Induction
****
GFP-Sham
HNF4a-Sham
GFP-PHx80%
HNF4a-PHx80%
B
% HNF4a+Ki67+/HNF4a+ cells
C
% HNF4a+pHH3+/HNF4a+ cells
D
Mitotic nuclei figure per 20x field
PHx80%
E
Cyp2e1
Cyp1a2
Cps1
Otc
Glul
Cyp7a1
Bsep
Apob
Alb
Ttr
F2
F5
F12
**
1.0
0.5
0
-0.5
F
% pHH3+PAS+/pHH3+ hepatocytes
***
G
Survival rate (%)
POD

**Suppl. Fig. 3 HNF4α viral-mediated overexpression alone does not increase survival after a SFSS-setting hepatectomy.**

(A) HNF4α gene expression in GFP-Sham (n=7) or HNF4α-Sham (n=8) 2 weeks after AAV8-TBG-HNF4α injection. **(B)** Quantification of the percentage of Ki67 positive hepatocytes in GFP-PHx80% (n=7), HNF4α-PHx80% (n=3) on POD3 in C57Bl/6J mice. **(C)** Quantification of the percentage of pHH3 positive hepatocytes in GFP-PHx80% (n=7), HNF4α-PHx80% (n=3) on POD3 in C57Bl/6J mice. **(D)** Quantification of mitotic features in 20x/field in GFP-PHx80% (n=7), HNF4α-PHx80% (n=3) on POD3 in C57Bl/6J mice. **(E)** Heatmap of genes expressions reflecting hepatocyte function on POD3. Z-score was calculated from DeltaCT and plotted on the heatmap. GFP-PHx80% (n=7), HNF4α-PHx80% (n=3). **(F)** Quantification of proliferating hepatocytes (pHH3%$^{+}$) stained with PAS on POD3 in GFP-PHx80% (n=6), HNF4α-PHx80% (n=3) and HNF4α/G-CSF-PHx80% (n=4). **(G)** Kaplan Meier survival curves in PHx80% (n=73), HNF4α-PHx80% (n=7) and HNF4α/G-CSF-PHx80%(n=8) in C57Bl6/J mice.

**Suppl. Fig. 4**

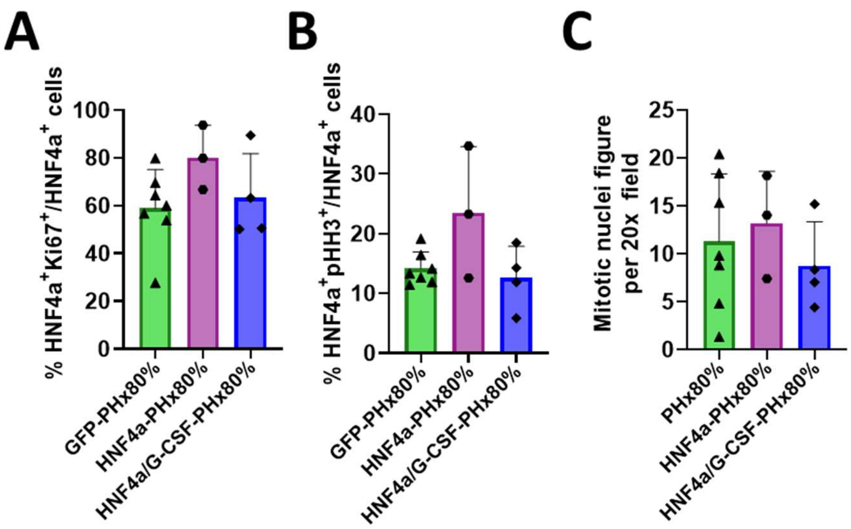


**Suppl. Fig. 4 HNF4α overexpression combined or not with G-CSF does not influence hepatocyte proliferation during regeneration.**

**(A)** Quantification of the percentage of Ki67 positive hepatocytes in GFP-PHx80% (n=7), HNF4α-PHx80% (n=3) and G-CSF/HNF4α-PHx80% (n=4) on POD3 in C57Bl/6J mice. **(B)** Quantification of the percentage of pHH3 positive hepatocytes in GFP-PHx80% (n=7), HNF4α-PHx80% (n=3) and G-CSF/HNF4α-PHx80% (n=4) on POD3 in C57Bl/6J mice. **(C)** Quantification of mitotic features in 20x/field in GFP-PHx80% (n=7), HNF4α-PHx80% (n=3) and G-CSF/HNF4α-PHx80% (n=4) on POD3 in C57Bl/6J mice.

**Supplementary table 1 Primers used for the heatmap of functional genes of the hepatocytes.**

| | Forward primers | Reverse primers |
|---|---|---|
| ***Cyp2e1*** | CGTTGCCTTGCTTGTCTGGA | AAGAAAGGAATTGGGAAAGGTCC |
| ***Cyp1a2*** | AGTACATCTCCTTAGCCCCAG | GGTCCGGGTGGATTCTTCAG |
| ***Cps1*** | CACCAATTTCCAGGTGACCA | TACTGCTTTAGGCGGCCTTT |
| ***Otc*** | AGGGTCACACTTCTGTGGTTC | CAGAGAGCCATAGCATGTACTG |
| ***Glul*** | GCTGCAAGACCCGTACCCT | TTCCACTCAGGTAACTCTTCCACA |
| ***Cyp7a1*** | AGCAACTAAACAACCTGCCAGTACTA | GTCCGGATATTCAAGGATGCA |
| ***Bsep*** | CTGCCAAGGATGCTAATGCA | CGATGGCTACCCTTTGCTTCT |
| ***Apob*** | AAGCACCTCCGAAAGTACGTG | CTCCAGCTCTACCTTACAGTTGA |
| ***Alb*** | GCTGCGCTGAAGCCAATC | GCTGAAATTCAGCAAGCACTGT |
| ***Ttr*** | TGTTCCGATACTCTAATCTCCC | TATACCCCCTCCTTCCAACC |
| ***F2*** | CCGAAAGGGCAACCTAGAGC | GGCCCAGAACACGTCTGTG |
| ***F5*** | CGCAACTAAGGCAGTTCTATGT | GCTAGATCGTGGCTTTTCTTTCT |
| ***F12*** | ATGACGGCTCTGTTGTTCCTG | CGGTGGTACTGAAAGGGAAAATG |
